# Functionality of BRCA1 supports the survival of prostate cancer cells during the development of castration resistance

**DOI:** 10.1101/2024.06.19.599365

**Authors:** Saiganesh Sriraman, Verneri Virtanen, Antti Kukkula, Mervi Toriseva, Anne Rokka, Anni Lumiainen, Johanna K. Ahlskog, Gun West, Matti Poutanen, Pekka Taimen, Maria Sundvall

## Abstract

Androgen deprivation therapy (ADT; castration) is the main treatment option for metastatic prostate cancer (PCa), but eventually, castration-resistant prostate cancer (CRPC) develops with no curative treatments. In CRPC, more than 20% of men carry mutations in DNA damage response (DDR) genes, including BRCA1/2. In this study, we elucidated the prostate tissue-specific functional role of BRCA1 protein. Our results indicate that DDR is dynamically regulated by androgen receptor (AR) signaling, and AR activation by the natural ligand dihydrotestosterone strongly downregulates the expression of BRCA1 in multiple cell lines. Consistent with these findings, our analyses of patient samples and mouse xenografts showed that DNA damage and BRCA1 expression were sustained after ADT. With unbiased mass spectrometry and bioinformatics approaches as well as experimentally, we found that BRCA1 interacts with Raptor, an mTORC1 component, and regulates the mTOR signaling pathway and PCa growth *in vitro*. Furthermore, we found that mTOR inhibition reduced the recruitment of DDR proteins, BRCA1 and Rad51, to DNA damage sites, creating a vulnerability towards DNA damage-inducing androgen deprivation. Moreover, we observed that BRCA1 supported ADT-induced activation of the oxidative stress sensor NRF2. Our findings shed further light on the complex DDR–AR interplay in PCa and suggest that, during PCa progression, BRCA1 expression may be retained due to the beneficial modulation of mTORC1 signaling in the AR environment by BRCA1.

**Significance:** Androgen receptor activation acts as a strong suppressor of BRCA1. Consequently, androgen deprivation activates BRCA1, which in turn promotes survival in castration resistance by supporting mTOR signaling and NRF2-mediated antioxidant processes.

## Introduction

Prostate cancer (PCa) is a prevalent cancer with significant morbidity and mortality worldwide [1]. Although treatment options for PCa have advanced over the years, metastatic PCa often eventually develops into castration resistance after the key growth driver androgen receptor (AR) signaling, is targeted and remains incurable [2, 3]. Castration-resistant prostate cancer (CRPC) has germline or somatic DNA damage response (DDR) gene alterations in approximately one-fourth of the cases [4]. DDR alterations leading to homologous recombination (HR) deficiency are considered to make cancer cells susceptible to poly ADP-ribose polymerase inhibitors (PARPi) by synthetic lethality. In response, several phase three clinical trials have led to the approvals of PARPi as monotherapy and in combination with antiandrogens for use in PCa in the presence of specified DDR alterations [5–8]. Despite the apparent importance of DDR in CRPC, relatively little is known about the tissue-specific functions of individual DDR proteins in PCa, although genes such as *BRCA1* and *BRCA2* have been widely studied in breast and ovarian cancer.

Previous preclinical studies have demonstrated an interplay between AR signaling and DDR in PCa [9–12]. Irradiation-induced DNA damage activates AR to localize into the nucleus and bind DNA in order to regulate transcription [9]. AR activation effectively reduces the rate of DNA damage by promoting DDR via non-homologous end joining (NHEJ) [9]. When NHEJ is inhibited, AR activation increases Rad51 foci formation in response to ionizing radiation, suggesting the involvement of enhanced HR [9]. Treatment with the AR inhibitor enzalutamide and knockdown of AR, in turn, have been suggested to reduce HR rate and susceptibility to PARPi [10, 11]. However, treatment with another AR inhibitor, apalutamide, has only suggested reduced NHEJ, whereas HR seems to remain functional [12]. The expression of several NHEJ and HR DDR pathway genes, as well as PARP1, has been suggested to be regulated by AR [9–12]. Together, these studies suggest a strong potential for the induction of AR signaling to enhance DDR via the regulation of PARP1, NHEJ, and HR. However, different studies have emphasized the regulation of distinct components, and it remains unclear whether these components contribute equally to achieving enhanced DDR. Moreover, a recent study questioned some of the previous conclusions showing that AR does not directly regulate DDR, and clinical studies have shown limited efficacy of PARPi plus ARi in HR-proficient patients without DDR mutations [13].

Interestingly, while BRCA1 mutations are the most common DDR alterations in both breast and ovarian cancers, BRCA1 mutations are relatively rare in PCa [14]. Furthermore, previous IHC studies using BRCA1 antibody (MS110) have reported a 10%–15% prevalence of high BRCA1 expressing PCa, while the expression in normal prostate is low and linked high BRCA1 expression to aneuploidy, proliferative state, and metastasis [15, 16]. Moreover, BRCA1 amplifications are observed in up to 2.9% of metastatic patients [17]. Thus, in PCa not only BRCA1 mutations but also retained high expression of BRCA1 protein plays a significant role in disease progression.

Therefore, in this study, we characterized the tissue-specific role of BRCA1 protein and the regulation of BRCA1 and DDR by AR signaling in PCa using cell lines, patient samples, and a mouse xenograft model of androgen deprivation therapy (ADT). We show that BRCA1 expression is retained and upregulated in response to ADT and conversely strongly repressed upon AR activation by its natural ligand, dihydrotestosterone (DHT). We show that AR inhibitor enzalutamide reduces BRCA1 expression as previously reported, but the enzalutamide effect is also present in AR-negative cells, suggesting the involvement of an AR-independent mechanism. Using bioinformatics and mass spectrometry analyses, we identify novel functional BRCA1 interactions with growth regulatory pathways in PCa. Moreover, reduction in BRCA1 levels impaired the growth of 3D PCa spheroids, supporting its role in growth regulation. Together, these results demonstrate a novel connection between BRCA1 and mTOR signaling in PCa and show that BRCA1 is regulated by AR signaling. In addition to having tumor-suppressive functions when mutated, the functional BRCA1 protein also supports oncogenic processes via tissue-specific interactions in PCa.

## Materials and methods

### In silico analysis of BRCA1 positively correlated genes using PCa patient datasets

Publicly available PCa datasets from cBioPortal (https://www.cbioportal.org/) were utilized to analyze genes positively correlated with *BRCA1* in PCa. The top 1500 correlated genes (FDR<0.05) from the datasets [4, 17–21] were used to identify 155 common BRCA1 co-expressed genes. This set of 155 genes was used as input gene identifiers in the Molecular Signatures Database (MSigDB, https://www.gsea-msigdb.org/gsea/msigdb/) to identify their association with hallmark gene sets and Gene Ontology: Biological Processes (GO:BP) gene sets. The results are represented as bar graphs denoted with significant p-values. The effect of R1881 on *BRCA1* and prostate-specific antigen (*PSA*) mRNA expression in LNCaP cells was analyzed using the GSE50936 dataset [22]. Expression data (log2) were downloaded from the R2: Genomics Analysis and Visualization Platform (http://r2.amc.nl).

### Cell culture and treatments

PC3, DU145, VCaP, LNCaP, and 22Rv1 cells (ATCC, Manassas, VA, USA) were cultured as described previously [23]. All cell lines were cultured in the presence of 1% penicillin-streptomycin solution (Sigma-Aldrich, St. Louis, MO, USA) except during the experiments, and were routinely tested to be mycoplasma-free and checked for authenticity.

Cells were seeded in 6-well plates for experiments other than immunofluorescence microscopy (24-well plates), IncuCyte, and 3D (transfected in 12-well plates and transferred to 96-well plates) assays. Androgen deprivation (AD) was performed using phenol-red free media containing 5% charcoal-stripped FBS (Gibco, Waltham, MA, USA; AD media, ADM). Glucose deprivation was performed by incubating the cells in media containing 5 mM glucose for 4 h prior to lysis, which was performed 48 h after transfection. Glutamine deprivation was performed using medium without glutamine for 2 or 4 h preceding lysis, which was performed 48 h after transfection. Serum starvation was performed using media without FBS for 24 h prior to lysis. Normal culture conditions were used as controls for deprivation experiments and are referred to as control media (CM).

DHT (Sigma-Aldrich) and R1881 (Sigma-Aldrich) were used to stimulate AR by incubating for 24 h prior to lysis and after incubation in ADM for 48 h unless specified otherwise. Treatments with AR inhibitor enzalutamide (Selleckchem, Houston, TX, USA) were performed for 24 h prior to lysis. mTOR inhibition was performed by incubating the cells with PP242 (Abmole, Houston, TX, USA) for 24 h prior to lysis. Cisplatin (Abmole) was used to induce DNA damage by incubating cells for 24 h prior to lysis. The proteasome inhibitor MG132 (Sigma-Aldrich) was used as a positive control to induce NRF2 expression by incubating cells for 24 h prior to lysis. Treatment groups compared were seeded simultaneously and lysed together.

### RNA interference and transfection

Transfection was performed 24 h after seeding the cells into the wells using the DharmaFECT Transfection Reagent (Dharmacon, Lafayette, CO, USA) and *Silencer*^®^ Select siRNA oligonucleotides (Ambion, Austin, TX, USA) listed in the Supplementary Table. The transfection solutions were prepared in Opti-MEM (Gibco) using 2.5 nM siRNA and 0.125% Dharmafect reagent 2 for PC3 cells, 10 nM siRNA and 0.5% Dharmafect reagent 3 for VCaP cells, and 5 nM siRNA and 0.125% Dharmafect reagent 3 for LNCaP cells. The transfection medium was replaced with regular media 24 h later to avoid excessive cell stress, and the treatments were applied. Knockdown efficiency was determined by Western blotting 48 h after transfection or at the end of the experiment.

### Proliferation assay

PC3 cells were subjected to BRCA1 knockdown using siRNA transfection, followed by trypsinization after 24 h and were plated in triplicate in 96-well plates. Cell proliferation (confluence) in a 2D culture was monitored in real time for up to 72 h using IncuCyte S3 (Essen Bioscience, Ann Arbor, MI, USA).

### Organotypic 3D growth assays

PC3 and LNCaP cells were grown in 3D using a previously published protocol [23]. The multispheroid average area was measured using 10 wells from each of the BRCA1 (siBRCA1-2/ siBRCA1-3) and control siRNA (siNeg1 and siNeg2) –treated cultures, using the IncuCyte S3 multispheroid analysis software. The cell viability in the IncuCyte assay was determined with the Cell Counting Kit-8 Assay (WST-8; Dojindo Molecular Technologies, Japan) and the absorbance were measured at 450 nm using a Wallac Victor2 1420 Multilabel Counter (PerkinElmer, Waltham, MA, USA).

### Cell lysis and immunoblotting

Cells were lysed and Western blotting was performed as previously described [23]. Equal amounts of 30–100 μg of total protein from the lysate supernatants were analyzed in the experiments. The list of antibodies used can be found in the Supplementary Table.

### Quantitative real-time PCR

RNA was isolated from VCaP and LNCaP cells post AD with or without DHT treatments with TRIsure (Meridian Bioscience, Cincinnati, OH, USA) or TRIzol reagent (Invitrogen, Thermo Fisher Scientific, Waltham, MA, USA), according to the manufacturer’s protocol. RNA was further purified using the RNA Clean & Concentrator-25 kit (Zymo Research, Irvine, CA, USA). The quantity and purity of the isolated RNAs were measured at 260 nm using a NanoDrop 2000 spectrophotometer (Thermo Fisher Scientific). cDNA was synthesized from 1 µg of total RNA, using the SensiFAST cDNA Synthesis Kit (Meridian Bioscience), following the manufacturer’s protocol. Reverse transcription reactions were performed in an Applied Biosystems Veriti™ Thermal Cycler (Thermo Fisher Scientific) as follows: 10 min at 25 °C for primer annealing, followed by 42 °C for 15 min for reverse transcription, 85 °C for 5 min for enzyme inactivation, and 4°C hold. Quantitative real-time PCR was performed in 96-well plates (Bio-Rad, Hercules, CA, USA) using 100 ng of cDNA (from each condition) and 10 µl TaqMan™ Fast Advanced Master Mix (Thermo Fisher Scientific), along with 1 µl respective TaqMan™ Gene Expression Assay (FAM) probes (Array ID (a) Hs02576345_m1, KLK3, FAM-MGB / 20X, (b) Hs03023880_g1, ACTB, FAM-MGB / 20X and (c) Hs01556193_m1, BRCA1, FAM-MGB / 20X), and nuclease-free water was added to reach a final reaction volume of 20 µl. The reaction was performed using the Bio-Rad CFX96™ Real-Time PCR System (Bio-Rad) with the following conditions; uracil-N-glycosylase activation at 50°C for 120 sec, followed by polymerase activation at 95°C for 20 sec, and lastly by 40 PCR cycles (denaturing for 3 sec at 95°C and annealing at 60°C for 30 sec), with real-time detection of the target sequence during the annealing phase. Gene expression was analyzed using the delta-delta Ct method.

### Immunofluorescence staining

Immunofluorescence analysis was performed according to a previously published protocol [23]. All the antibodies used are listed in the Supplementary Table. VCaP cells were subjected to vehicle (DMSO) or cisplatin treatment with or without the pan-mTOR inhibitor PP242 for 24 h prior to fixation. The cells were imaged using a 3i CSU-W1 Spinning disc confocal microscope (Zeiss, Oberkochen, Germany) with a 63X oil objective.

### Prepraring the nuclear and cytoplasmic fractions

VCaP cells subjected to treatments were harvested with trypsin and washed with cold PBS, and 5 million cells from each condition were used for fractionation. NE-PER™ Nuclear and Cytoplasmic Extraction Reagents (Thermo Fisher Scientific) supplemented with 2% cOmplete EDTA-free protease inhibitor cocktail (Roche, Basel, Switzerland), 10 mM Na_4_P_2_O_7_, and 1 mM Na_3_VO_4_ phosphatase were added to the cells just prior to the lysis, and extraction was performed according to the manufacturer’s instructions. Equal amounts of total protein from the cytoplasmic and nuclear fractions were analyzed by immunoblotting against MEK-1 (a cytoplasmic marker) and Lamin B1 (a nuclear marker). Information on the antibodies is provided in the Supplementary Table.

### Immunoprecipitation of BRCA1 interacting proteins

PC3 cells were grown in multiple 10 cm dishes for 24 h after plating before lysis or 48 h after transfection, followed by lysis with a buffer containing reduced NaCl (100 mM) along with regular protease and phosphatase inhibitors. After lysis, 3–4 mg of protein was used for immunoprecipitation (IP). After pre-clearance with G-Sepharose beads (GE Healthcare, Chicago, IL, USA), 2 µg of BRCA1 or IgG antibody were added and incubated at 4°C overnight. G-Sepharose beads were added to lysates for 2 h incubation at 4°C. Beads were washed, and samples were boiled in 6X SDS and run on single or duplicate gels. See the Supplementary Table for antibody information.

For mass spectrometry analysis, PC3 cells (30 x 10 cm plates) and LNCaP cells (40 x 10 cm plates) were lysed and precleared with G-Sepharose beads for 1 h at 4°C. Then 0.5 µg of BRCA1 or IgG control antibody was added per mg of sample protein and incubated overnight at 4°C. Samples were incubated with G-Sepharose beads and washed five times with lysis buffer containing 150 mM NaCl before elution three times in 100 mM glycine pH 2.5, followed by elution in Tris-HCl pH 8. Samples were concentrated using Amicon filters (Millipore, Burlington, MA, USA) with a 10 kDa cutoff and 6X SDS was added before boiling the samples. Samples were run into gels and gel pieces were extracted by cutting above the IgG band at 50 kDa after PageBlue staining.

### LC-ESI-MS/MS Analysis

Immunoprecipitated BRCA1 and IgG control samples separated on SDS-PAGE gel were cut into several fractions before in-gel trypsin digestion. The digested peptides were dissolved in 0.1% formic acid and analyzed by mass spectrometry. LC-ESI-MS/MS analyses were performed on a nanoflow HPLC system (Easy-nLC1000) coupled to a Q Exactive mass spectrometer (Thermo Fisher Scientific) equipped with a nano-electrospray ionization source. Peptides were first loaded on a trapping column (100 µm ID x 2 cm) and subsequently separated inline on and analytical column (75 μm ID x 15 cm). The packing material for the both columns was ReproSil-Pur 5 μm 200 Å C18-AQ (Dr. Maisch HPLC GmbH, Ammerbuch, Germany).

The mobile phase consisted of water with 0.1% formic acid (solvent A) or acetonitrile/water (80:20 (v/v)) with 0.1% formic acid (solvent B). A 20 min gradient from 8 to 43% B was used to elute the peptides.

MS data were acquired by an information-dependent acquisition method consisting of an Orbitrap MS survey scan of a mass range 300-2000 m/z followed by HCD fragmentation for the 10 most intense peptide ions.

Data files were searched for protein identification using Proteome Discoverer 2.2 software connected to an in-house server running Mascot software (Matrix Science, Chicago, IL, USA). Data were searched against the SwissProt database with a taxonomy filter ‘homo sapiens’. Samples were merged together in data processing in order to represent the peptide content of the whole digested gel piece. At least two peptides with medium or high confidence, were required per protein.

### Murine VCaP xenografts of early ADT response

The VCaP xenograft sections were generated according to previously described protocol and permissions [24]. The subcutaneous tumors were grown for 4 weeks and surgical castration was performed. The mice were sacrificed either two days (Cas 2D) or five days (Cas 5D) after castration. The Intact group, serving as the non-castrated control, was sacrificed five days after the surgery on the treatment groups. Tumors generated under these conditions were collected and fixed in formalin, followed by sectioning and sample preparation.

### Immunohistochemistry, digital image analyses, and ethical permissions

Tissue sections from patient samples in a tissue microarray and from tumor xenografts were deparaffinized with xylene (3 x 7 min) and rehydrated in absolute ethanol (2 x 2 min) and 96% ethanol (2 x 2 min), followed by three washes with distilled water. Antigen retrieval was performed in citrate buffer (pH 6) using a microwave for 7 min at 600 W, 7 min at 450 W and the sections were left to cool down for 20 min. For pBRCA1 and γH2Ax staining, antigen retrieval was performed using a decloaking chamber (Biocare Medical NxGen, Pacheco, CA, USA) for 20 min. After three washes with distilled water, sections were blocked with 3% hydrogen peroxidase for 10 min at RT, washed with Tris-HCl buffer, followed by pre-protein block in antibody diluent for 10 min at RT, and incubated using antibodies and concentrations specified in the supplemental table for 60 min at RT. After washing with Tris-HCl buffer, the sections were incubated with secondary antibody for 30 min at RT and washed again with Tris-HCl buffer. DAB incubation was washed with distilled water after 10 min at RT, and the sections were counterstained with Mayer’s hematoxylin for 1 min at RT, followed by washing with distilled water and dehydration twice in 96% ethanol, twice in absolute ethanol, and three times in xylene.

The use of patient tissue material was approved by the Ethics Committee of the Hospital District of Helsinki and Uusimaa (84/13/03/00/2014; §3 30.01.2015), the Hospital District of Southwest Finland (number T206/2014), and the National Supervisory Authority for Welfare and Health (VALVIRA, 8008/06.01.03/2014).

The slides were scanned using a Pannoramic P1000 slide scanner (3DHISTECH, Budapest, Hungary). Xenograft whole tumor sections were measured for total tumor area in QuPath version 0.4.4 using a pixel classifier after refining the area analyzed using manual segmentation. The areas of strong staining were detected using the QuPath pixel classifier. For the patient samples, two researchers independently performed manual scoring.

### Statistical analysis

Statistical analyses were performed using R Studio (R Project for Statistical Computing) and GraphPad Prism 10.1.2 software. Data extracted from cBioPortal were generated using the default methods of the v4.1.9. Spearman’s rank correlation coefficient was used to analyze correlations. Two-tailed unpaired Student’s t-test or Mann-Whitney U test were used for comparisons between two groups. Analysis of Variance (ANOVA) test was used to analyze the difference between the means of more than two groups.

### Data availability

Raw data generated in this study are available upon reasonable request from the corresponding author.

## Results

### AR mediates transcriptional downregulation of BRCA1 expression

Since activation of AR has been suggested to regulate HR and inhibition of AR has been shown to downregulate BRCA1 expression, we wanted to study whether BRCA1 protein was induced by activated AR. In this study, we used VCaP (AR-amplified, TP53-mutated, TMPRSS2-ERG fusion), LNCaP (AR-mutated, CHEK2-SNV, PTEN loss), 22Rv1 (AR splice variant, BRCA2-mutated, ATM-SNV, TP53-mutated, TMPRSS2-ERG fusion), PC3 (AR-negative, PTEN loss), and DU145 (AR-negative, TP53-mutated, Rad50-SNV, BRCA2-SNV) PCa cell lines [25]. To study BRCA1 protein levels in response to AR activation, we treated the highly AR-responsive VCaP cell line with different DHT concentrations (0.05 to 10 nM) for 24, 48, and 72 h after AD for 72, 48, and 24 h, respectively. We utilized androgen-depleted samples and cells grown under normal culture conditions as controls. We observed increased expression of the AR transcriptional targets PSA and FKBP5 in the treatment range of 0.5–10 nM (Figure 1A and B).

**Figure 1.**
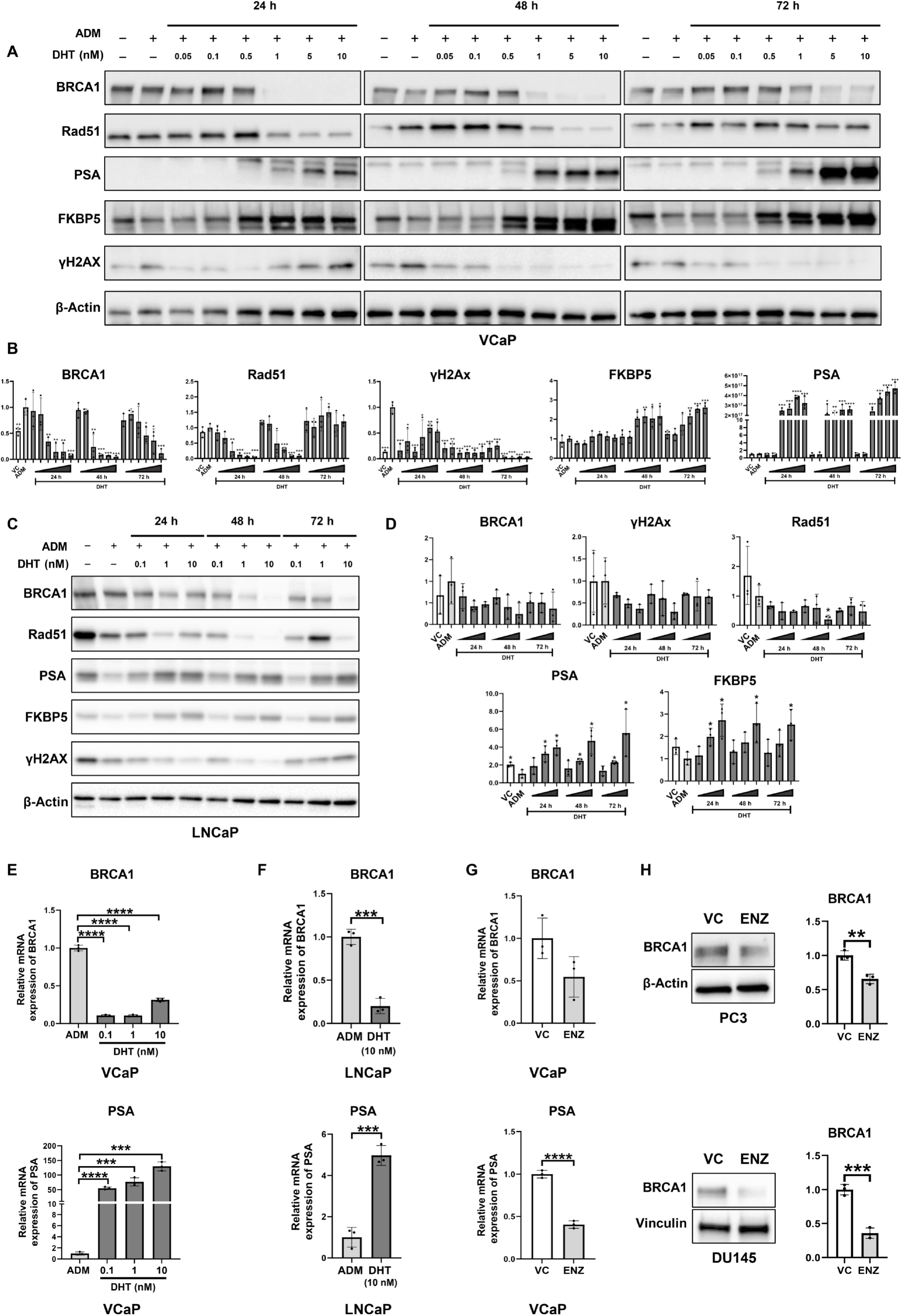
BRCA1 and Rad51 expression is repressed by AR activation. (A) Representative Western blots of BRCA1, Rad51, PSA, FKBP5, γH2Ax, and loading control expression in VCaP cells treated with DHT (0.05–10 nM) for 24, 48, and 72 h after incubation in ADM starting 96 h before lysis with vehicle (methanol) and ADM (96 h) as controls. (B) Bar graphs depicting pooled Western blot densitometry results of 3 independent biological repeats in VCaP cells. (C) Representative Western blots of LNCaP cells treated with DHT (0.1–10 nM) for 24, 48, and 72 h after incubation in ADM starting 96 h before lysis with vehicle (methanol) and ADM (96 h) as controls. (D) Bar graphs depicting pooled Western blot densitometry results of 3 independent biological repeats in LNCaP cells. (E) Bar graphs depicting relative expression of *BRCA1* and *PSA* mRNA by real-time qPCR in VCaP cells subjected to ADM for 72 h or DHT (0.1 nM to 10 nM) for 24 h after 48 h of ADM. (F) Bar graphs depicting relative expression of *BRCA1* and *PSA* mRNA by real-time qPCR in LNCaP cells subjected to ADM for 72 h or DHT (10 nM) for 24 h after 48 h of ADM. (G) Bar graphs depicting relative expression of *BRCA1* and *PSA* mRNA by real-time qPCR in VCaP cells treated with vehicle (DMSO) or enzalutamide (1 µM) for 24 h. (H) Representative Western blots and bar graphs of pooled BRCA1 expression from Western blot densitometry against loading control in PC3 cells treated with vehicle (DMSO) or enzalutamide (1 µM) for 24 h. (I) Representative Western blots and bar graphs of pooled BRCA1 expression from Western blot densitometry against loading control in DU145 cells treated with vehicle (DMSO) or enzalutamide (1 µM) for 24 h. The mean and standard deviation (SD) from three experiments are represented as bar graphs (*p < 0.05, **p < 0.01, ***p < 0.001, and ****p < 0.0001 as determined by *t-test*).

In contrast, with the same DHT treatment concentrations BRCA1 protein was downregulated in a dose-dependent manner most severely at 24- and 48-hour time points. As a readout for DNA damage, we used γH2Ax, which was strongly downregulated already with 0.05 nM DHT. Another essential functional HR protein, Rad51, was downregulated in a manner similar to that of BRCA1. BRCA1 downregulation was also observed in response to the synthetic androgen R1881 (Supplementary Figure S1A). These results suggested that activated AR was unexpectedly downregulating rather than inducing BRCA1 expression in the AR-amplified VCaP cell line. Moreover, our results using γH2Ax as a readout showed that AR stimulation efficiently reduced DNA damage, despite concurrently reduced key HR protein levels, suggesting that AR stimulation, already at low doses, either proficiently arrested the initiation of DNA damage or possibly enhanced alternative repair.

Stimulation of AR signaling was performed similarly in LNCaP cells (Figure 1C and D). As expected, PSA and FKBP5 responses were less extreme, yet highly dose-dependent in comparison to the VCaP cell line which yielded a stronger response in AR target gene expression. We observed trends matching the VCaP cell line results for BRCA1, Rad51, and DNA damage readout, although in LNCaP cells downregulation was less severe, and only Rad51 reached significance. Taken together, our results implied that the HR proteins BRCA1 and Rad51 were not upregulated in a manner similar to that of PSA and FKBP5 after AR stimulation.

We also examined the correlation between *BRCA1* and AR transcriptional targets in PCa patient and patient-derived organoid datasets (Supplementary Figure S1B and C). The datasets showed a trend of negative correlation, providing further evidence for the downregulation function of AR over BRCA1. Although AR signaling strongly regulated BRCA1 expression, no reciprocal regulatory effect on AR activity was observed during BRCA1 knockdown (Supplementary figure S1D).

Next, we wanted to determine how the activated AR mediated BRCA1 downregulation. Since BRCA1 seemed to be regulated at similar doses and time points as the transcriptional targets, BRCA1 could also be regulated by AR at the transcriptional level. To test this, we performed qRT-PCR in both VCaP (0 to 10 nM) and LNCaP (0 and 10 nM) cells and observed downregulated BRCA1 mRNA levels in response to AR stimulation by DHT (Figure 1E and F). A similar response to R1881 was observed in a published RNA dataset (Supplementary Figure S1E) [22]. This suggested that, upon activation, AR mediated repression of *BRCA1* transcription.

As AR stimulation by DHT decreased BRCA1 levels, it was expected that the inhibition of AR activation would reciprocally increase BRCA1 expression. However, previous reports have shown that BRCA1 was downregulated in response to AR antagonist enzalutamide in VCaP and LNCaP cell lines [10, 11]. Indeed, we observed a similar downregulation of *BRCA1* with enzalutamide (Figure 1E–G). Thus, we tested whether the enzalutamide-induced downregulation was truly AR-mediated. We treated AR-negative PC3 and DU145 cells with enzalutamide and observed similar downregulation (Figure 1H). This demonstrated that enzalutamide could downregulate BRCA1 expression also in the absence of AR, suggesting AR-independent non-target effects.

### AD induces DNA damage in PCa cells

Considering that activated AR downregulates BRCA1 and that downregulation by enzalutamide does not require AR expression, we wanted to clarify how the reduced activation of AR would affect BRCA1 expression. We used media supplemented with steroid-free serum to mimic ADT by depriving AR of its ligand. We selected three heterogeneously AR-altered AR-positive cell lines to represent the different AR signaling states. VCaP cell line represented AR-amplified PCa, LNCaP cell line represented AR-mutated PCa, and 22Rv1 cell line represented PCa with high expression of a constitutively active AR-splice variant AR-V7. As expected, VCaP and LNCaP cells showed downregulation of PSA, while 22Rv1 retained PSA expression at 24 h (Figure 2A–D).

**Figure 2.**
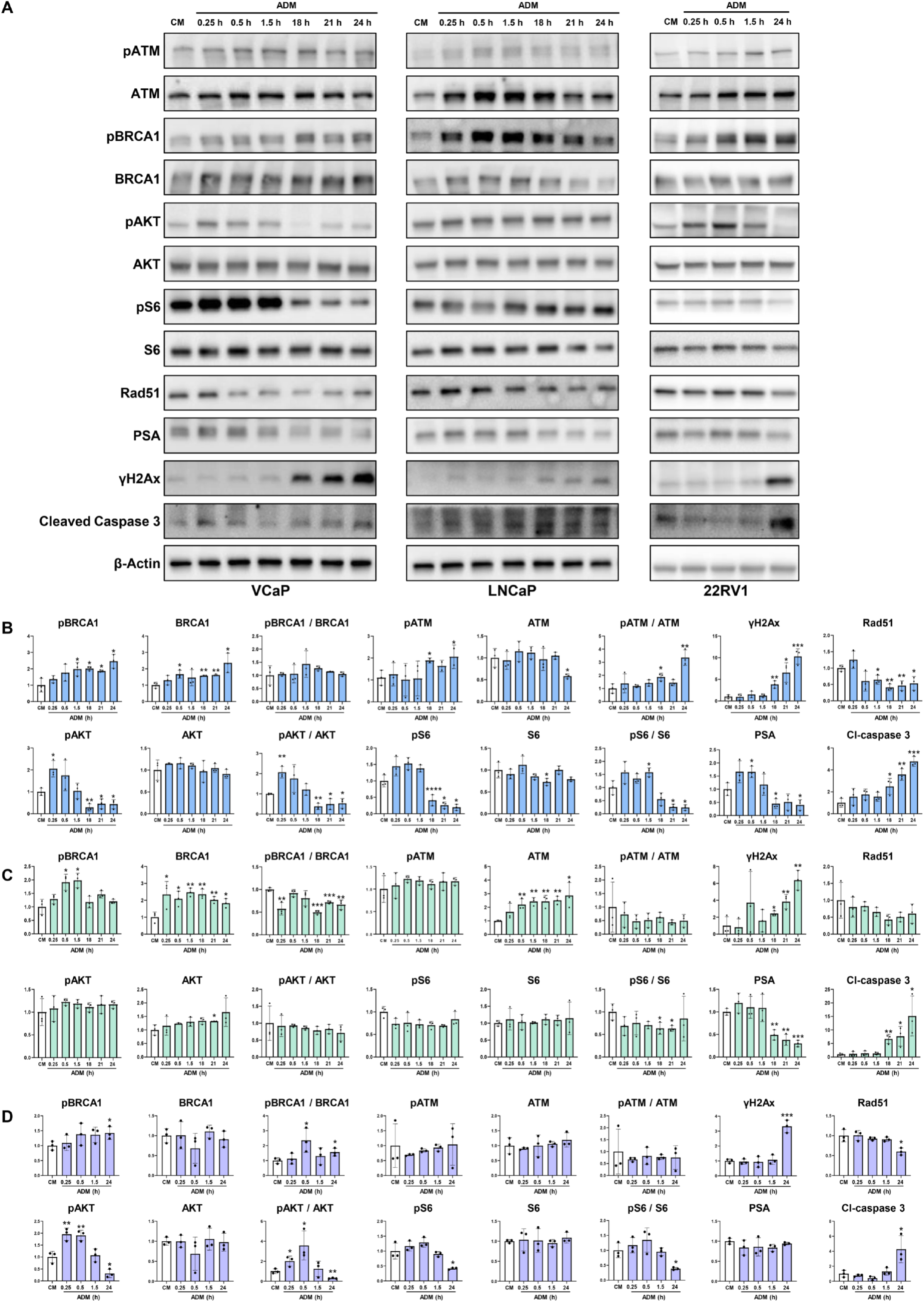
AD induces DNA damage and cell death in PCa. (A) Representative Western blots depicting expression of key HR components, AR targets, and markers of PI3K-Akt/mTOR signaling, DNA damage, and apoptosis with loading control expression in VCaP, LNCaP, and 22RV1 cells subjected to ADM (0.25–24 h) along with control (normal culture media). (B–D) Bar graphs depicting pooled Western blot densitometry results of 3 biological repeats. The mean and standard deviation (SD) from three experiments are represented as a bar graph (*p < 0.05, **p < 0.01, and ***p < 0.001 as determined by *t-test*).

We observed upregulation of total BRCA1 in VCaP and LNCaP cells, while for the constitutively AR-activated 22Rv1 cells with reduced PSA-responsiveness, BRCA1 levels were constant (Figure 2A–D). We also observed increased levels of phosphorylated BRCA1 at the ATM target phosphorylation site [26] in all cell lines suggesting the activation of BRCA1. However, in LNCaP cells, phosphorylation was only observed at shorter time points, whereas in other cell lines (PTEN-expressing, p53-mutated, and TMPRSS2-ERG fusion-positive cells), a significant increase was observed at later time points. This suggested that the BRCA1 protein was dynamically regulated in response to AD in different PCa cell lines.

Rad51 was observed to have a trend of downregulation in all cell lines, significant in VCaP and 22Rv1 cells (Figure 2A–D). This suggested that while *BRCA1* regulation might be explained by the presence or absence of repression mediated by activated AR, *RAD51* may have another layer of regulation in the AD context.

We observed upregulated γH2Ax levels in all cell lines, notably also in 22Rv1 cell line, where no PSA response was observed (Figure 2A–D). Similarly, cleaved caspase-3 was upregulated in all the cell lines. These data suggested that AD induced DNA damage and eventually apoptosis in PCa cell lines. Interestingly, the observed BRCA1 upregulation was thus insufficient to prevent DNA damage.

We also analyzed other DDR proteins after AD. VCaP cells showed an increase in pATM and no change in total ATM was observed (Figure 2A–D). In LNCaP cells, pATM showed no change and ATM expression increased over time. Furthermore, ATM-activated CHK2 phosphoprotein and total CHK2 were also studied along with PARP1 and CtIP in VCaP cells (Supplementary Figure S2A). These proteins were mainly unchanged except for a passing PARP1 downregulation. The lack of change in CHK2 and CtIP levels suggests incomplete initiation of double-strand break signaling, potentially signifying an impaired HR.

S6 and Akt, as readouts of the PI3K-Akt/mTOR signaling pathway involved in the regulation of cell growth, survival, and protein synthesis, showed an initial surge of phosphorylation in response to AD but were strongly dephosphorylated at later time points. This pattern was not present for Akt and barely detectable for S6 in LNCaP cells, in which Akt was constitutively active due to PTEN loss. Similar responses were observed in VCaP and 22Rv1 cells, although PSA regulation was unclear in 22Rv1 cells owing to retained AR-V7 activity. This might suggest that PI3K-Akt/mTOR signaling is associated with full-length AR activity, some of which is also present in 22Rv1 cells.

Taken together, our results demonstrated that the PI3K-Akt/mTOR pathway is activated shortly after AD, followed by strong induction of γH2Ax in all PCa cell lines tested in the presence of BRCA1 activation suggesting that BRCA1 is not the limiting factor for the DNA damage process in response to AD.

### Early time points of ADT induce DNA damage and BRCA1 phosphorylation in murine VCaP xenografts

To confirm our results *in vivo*, we performed ADT on murine VCaP xenografts by orchiectomy and sacrificed the mice two and five days after castration (Figure 3A). PSA and tumor volume were measured weekly and, as expected, ADT reduced both, suggesting that AR activity was successfully reduced and led to cell death (Supplementary Figure S3A and B). Additionally, tumor weight measured at sacrifice and area measured from sample slides showed a timepoint-dependent trend of decrease, providing further support for cell death expectedly occurring in our model (Supplementary Figure S3C and D).

**Figure 3.**
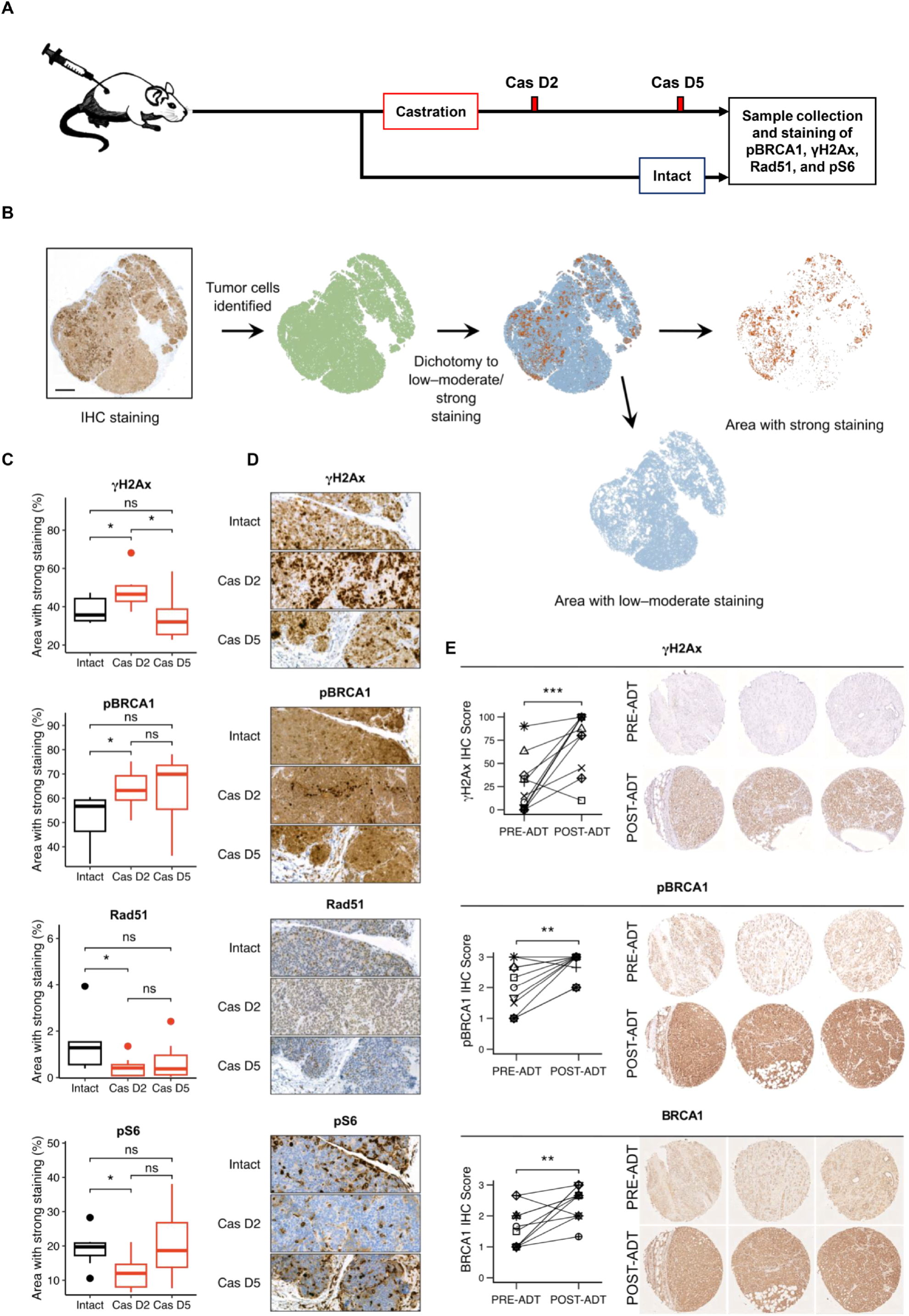
BRCA1 is phosphorylated and its expression is retained in ADT. (A) Schematic figure representing the sample generation process of Intact, Cas D2, and Cas D5 VCaP mice xenograft tumors. (B) A Schematic figure representing the analysis pipeline in QuPath for scoring the IHC staining. (C) Boxplots depicting the percentages of strong IHC staining of pBRCA1, Rad51, γH2Ax, and pS6 in VCaP xenograft tumors (ns = not significant and *p < 0.05 as determined by *Mann-Whitney U test*). (D) Representative images of IHC staining in VCaP xenograft tumor (E) Representative IHC staining and scoring of γH2Ax, pBRCA1 and BRCA1 from PCa samples obtained from the same patients before and after starting ADT (representative images shown, total n=11 patients, *p < 0.05 and **p < 0.01 as determined by *t-test*).

We performed IHC on xenograft tumors using γH2Ax, pBRCA1, Rad51, and pS6 antibodies. Scanned IHC samples were analyzed in the QuPath software using pixel classification to identify tumor cells and tumor areas with strong positive staining (Figure 3B). We observed changes in the staining of the VCaP xenografts from day two resembling our results seen in VCaP cells at the 24 h timepoint, indicating that these proteins were indeed regulated *in vivo* and, importantly, by the absence of androgen instead of the absence of other steroids removed from cell serum (Figure 3C and D). Samples from day five showed similar trends, albeit with greater variability, potentially suggesting that the increased DNA damage may have contributed to increased cell death rather than a continued accumulation of damage in surviving cells. Consequently, it is possible that the cells with the most severe DNA damage had already perished by day five, leading to the observed variability. Taken together, these *in vivo* data verified our *in vitro* findings, suggesting that AD induced DNA damage and increased BRCA1 activation.

### ADT induces expression and phosphorylation of BRCA1 and increases DNA damage in PCa patients

Samples were collected from 11 patients who underwent ADT treatment. Paired samples were collected before and after the ADT. 64% of post-ADT samples were of metastatic origin, and 64% of patients were considered castration-resistant at the time of post-ADT sample collection. Patient characteristics are presented as a table (Table 1). We performed IHC on patient samples using BRCA1, pBRCA1, and γH2Ax antibodies (Figure 3E). We scored the stained paired samples and observed an increase in the staining intensity for all three proteins. This suggested that DNA damage and cells expressing BRCA1 in an active phosphorylated state became enriched post-ADT and in metastases. Thus, activation of the BRCA1 protein and DNA damage may have potentially contributed to the development of resistance to ADT in these patients.

**Table 1.** Baseline characteristics of the patients. PSA, prostate-specific antigen; ADT, androgen deprivation therapy; RT, radiotherapy; TURP, transurethral resection of the prostate; CRPC, castration-resistant prostate cancer.

| Age (years) at initial diagnosis of prostate cancer |  |
| --- | --- |
| Median (range) | 58 (54–79) |
| Gleason score at initial diagnosis of prostate cancer (n, %) |  |
| 7 | 4 (36) |
| 8 | 3 (27) |
| 9 | 4 (36) |
| PSA (µg/L) level at primary diagnosis |  |
| Median (range) | 13 (3.9–57) |
| Initial treatment (n, %) |  |
| ADT + RT | 3 (27) |
| Surgery + ADT | 6 (54) |
| ADT only | 2 (18) |
| Time since start of ADT to post-ADT sample (months) |  |
| Median (range) | 66 (18–107) |
| Other hormonal treatments after ADT (n, %) |  |
| AR inhibitors | 11 (100) |
| Abiraterone | 1 (9) |
| Origin of post-ADT sample (n, %) |  |
| Prostate (TURP) | 4 (36) |
| Metastasis | 7 (64) |
| Clinical castration resistance at post-ADT sample (n, %) |  |
| CRPC | 7 (64) |
| No* | 4 (36) |
| Metastases** (n, %) |  |
| None | 1 (9) |
| Bone only | 2 (18) |
| Lymph node only | 1 (9) |
| Bone + lymph node or visceral only | 6 (54) |
| Visceral | 5 (45) |

\* Relapse after ADT but no CRPC at the time of post-ADT sample, \*\*at the time of post-ADT sample.

### Mass spectrometry identifies novel interactors of BRCA1 in PCa cells

The enriched BRCA1 expression found in post-ADT patient samples suggested that BRCA1 protein may have a tumor-promoting role in PCa progression. Therefore, we aimed to explore the tissue-specific functions of BRCA1 in PCa. We chose mass spectrometry of BRCA1-pulldown samples from LNCaP and PC3 cell lines as an unbiased method for exploring BRCA1 interactions in PCa (Figure 4A). Our analysis identified BRCA1 and well-characterized known interactors BRIP1, BARD1, and ATM, reassuring that the assay worked as expected [26–28]. In addition, we identified a considerable number of interactors predicted by the STRING database, including CASC3, CDH1, DDB1, DNMT1, HNRNPM, KNTC1, LMNA, PABPC1, UBA1, USP9X, and VCP (Supplementary Figure S4A).

**Figure 4.**
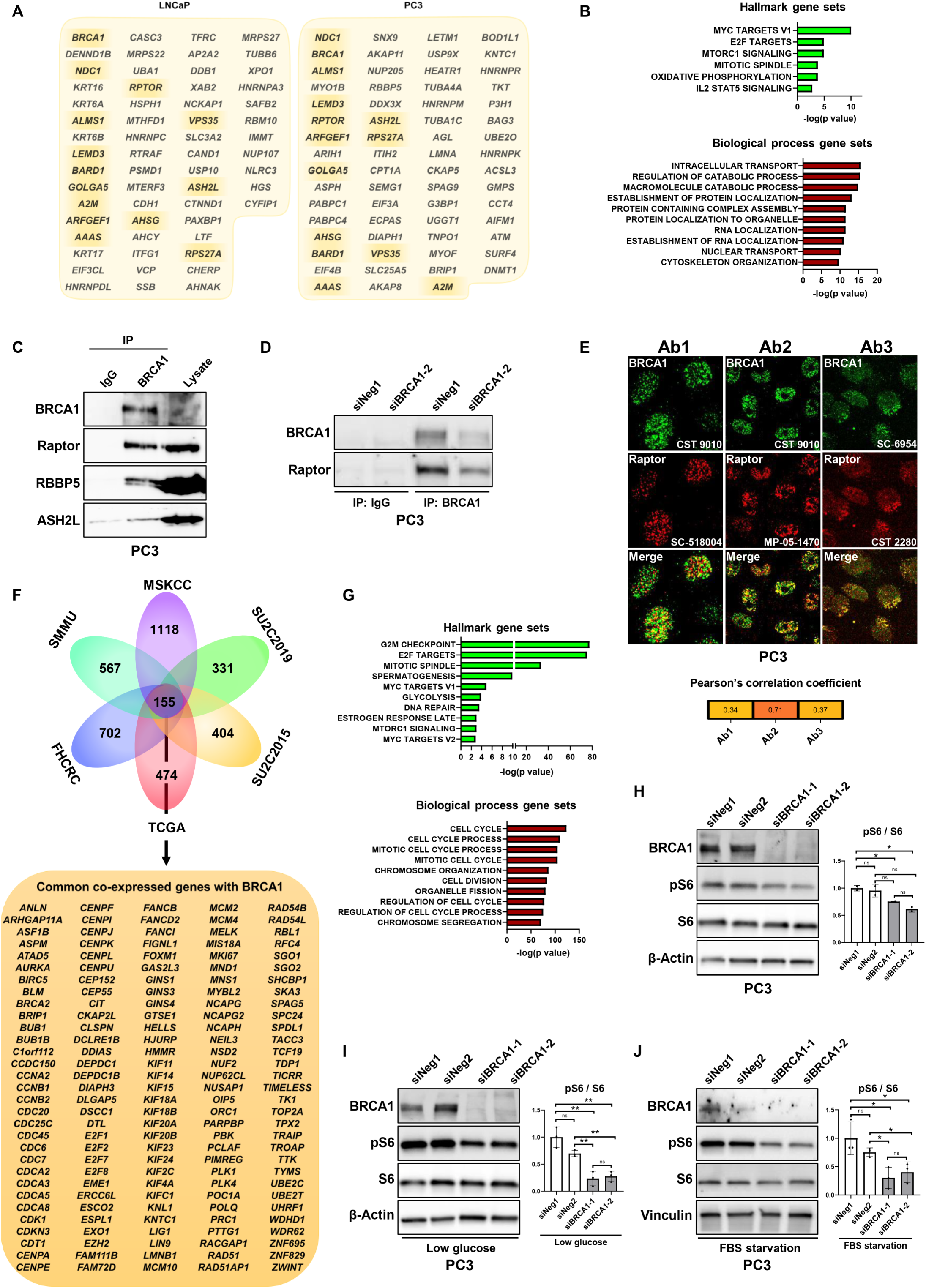
BRCA1 associates with key signaling pathways and regulates mTOR signaling in PCa. (A) Lists representing mass spectrometry hits from PC3 and LNCaP cell BRCA1-pulldown samples with common hits highlighted. (B) Significant associations of mass spectrometry hits with MSigDB hallmark gene sets and the top ten significant GO:BP gene sets. (C) Representative Western blot depicting BRCA1, Raptor, RBBP5, and ASH2L in IgG and BRCA1 antibody pulldowns in PC3 cell lysates. (D) Representative Western blot depicting BRCA1 and Raptor in IgG and BRCA1 antibody pulldowns of PC3 cell lysates with and without silencing BRCA1. (E) Immunofluorescence staining of BRCA1 (green) and Raptor (red) represented as both single and merged images with Pearson’s correlation coefficient depicting colocalization for signal from three combinations of antibodies in PC3 cells. (F) Venn diagram and list representing the top 155 mRNAs co-expressed with *BRCA1* consistently in six PCa patient datasets available on cBioPortal. (G) Top ten significant MSigDB hallmark gene sets and GO:BP gene sets that overlapped with the top 155 mRNAs concordantly co-expressed with *BRCA1* in PCa patients. (H–J) Western blot depicting BRCA1, pS6, and S6 with loading control in PC3 cells transfected with siNeg1, siNeg2, siBRCA1-1, or siBRCA1-2 48 h before lysis under normal culture conditions (n=2), glucose deprivation (n=3; 4 h before lysis), or serum deprivation (n=3; 24 h before lysis) conditions. The mean and standard deviation (SD) from three experiments are represented as bar graph unless otherwise indicated (ns = not significant, *p < 0.05, and **p < 0.01 as determined by *t-test*).

We further characterized the significant mass spectrometry hits by computing their overlaps with MSigDB GO:BP and Hallmark gene sets to explore the potential functions of BRCA1 interactors in PCa [29, 30]. Processes involved in catabolism, transport, protein and RNA localization, cytoskeleton organization, and pathways associated with MYC, E2F, mTORC1, mitotic spindle, oxidative phosphorylation, and IL2-STAT5 signaling were detected (Figure 4B).

### Bioinformatic analysis of BRCA1 in PCa patients

In a second unbiased approach to characterize the potential BRCA1 roles in PCa, we used mRNA data from publicly accessible datasets to detect co-expressed genes in patients. We combined the six datasets and identified the top 155 common genes that were positively correlated with *BRCA1* (Figure 4F). The identified co-expressed genes were investigated for their signature roles in different pathways and biological processes using MSigDB (Figure 4G). Pathways, such as the G2M checkpoint, E2F targets, glycolysis, DNA repair, estrogen response, MYC targets, mTORC1 signaling, and GO:BP categories, including cell cycle process and regulation, mitosis, chromosomal organization and segregation, cell division, and organelle fission, were detected. This supported the mass spectrometry findings of a potential connection between BRCA1 and mTOR in PCa. Together, these results suggested that *BRCA1* was involved in several essential functions related to cell division, development, and G2/M cell cycle processes, as well as important signaling pathways such as DNA repair, mTORC1, MYC, and glycolysis in PCa.

### Validation of mass spectrometry results using IP

To technically validate our mass spectrometry hits, we focused on selected potential novel interactors. We chose the mTOR adapter protein Raptor, which promotes mTORC1 signaling and activation of S6, as mTORC1 was indicated by both mass spectrometry and co-expression analysis. Additionally, the COMPASS complex components ASHL2 and RBBP5 were selected because these hits represented proteins from a complex that had previously been connected to PCa [31]. We performed Western blot for BRCA1-immunoprecipitation of PC3 cell line lysates and detected Raptor, ASHL2, and RBBP5 in the pulldown sample (Figure 4C).

By knocking down BRCA1 using siRNA, we observed reduced Raptor protein levels in the immunoprecipitation Western blot analysis (Figure 4D). Furthermore, we observed nuclear colocalization of BRCA1 and Raptor by immunofluorescence confocal microscopy employing two distinct antibodies detecting BRCA1 and three different antibodies against Raptor (Figure 1E). This suggested that BRCA1 had previously unreported interaction with Raptor and may have Raptor-mediated interplay with mTOR signaling in PCa cells.

### BRCA1 is functionally connected to mTOR signaling in PCa cells

To study the effect of BRCA1 on mTORC1 signaling, we silenced BRCA1 in PC3 cells and observed a trend of decreased pS6 expression (Figure 4H). The downregulation became significant when the cells were challenged with low glucose, FBS starvation, or low glutamine (Figure 4I and J, and Supplementary Figure S4B). Consistent with these results, increased phosphorylation of S6 was observed in PC3 cells stably transfected with the HA-BRCA1 construct to overexpress BRCA1 protein when compared to control cells expressing HA-tag construct without insert (Supplementary figure S4C). This suggested that in AR-independent castration-resistant cells, under low-nutrient conditions, the activation of the mTORC1-mediated protein synthesis pathway is regulated by the availability of BRCA1.

### BRCA1 supports PCa cell spheroidal growth

Although BRCA1 is considered a tumor suppressor, our results suggested that in PCa cells, lowering BRCA1 levels downregulated the activation of key survival and growth signaling via the mTORC1 pathway. Therefore, we tested whether BRCA1 knockdown affected PCa cell growth. We chose to use siRNA as our method of gene silencing, given its ability to retain protein expression partially, thereby protecting cell viability from the complete loss of essential functions of BRCA1. We found that *BRCA1* knockdown did not alter the confluence of PC3 cells during three days of growth, suggesting that 2D plastic adherent cell culture viability was unaffected by BRCA1 knockdown (Figure 5A). To test whether the unaffected state was limited to a 2D setting, we performed a 3D organotypic cell culture in the basal membrane extract with *BRCA1* knockdown. We observed that in PC3 cells, the BRCA1 knockdown spheroids remained significantly smaller compared to controls over the seven-day culture period, and growth inhibition was evident already at day four (Figure 5B and C). The WST-8 assay suggested that the overall viability was significantly reduced upon *BRCA1* knockdown (Supplementary Figure S4D). Western blotting was used to verify the knockdown, and pooled data from three experiments showed a similar reduction in spheroid size for both AR-expressing LNCaP cells and AR-negative PC3 cells (Figure 5D–H). Our data demonstrated that while 2D culture was unaffected by knockdown of BRCA1 until three days, cells growing in a 3D environment showed reduced spheroid size starting from four days. Taken together, these results suggested that BRCA1 function in PCa became a limiting factor for growth in a 3D environment.

**Figure 5.**
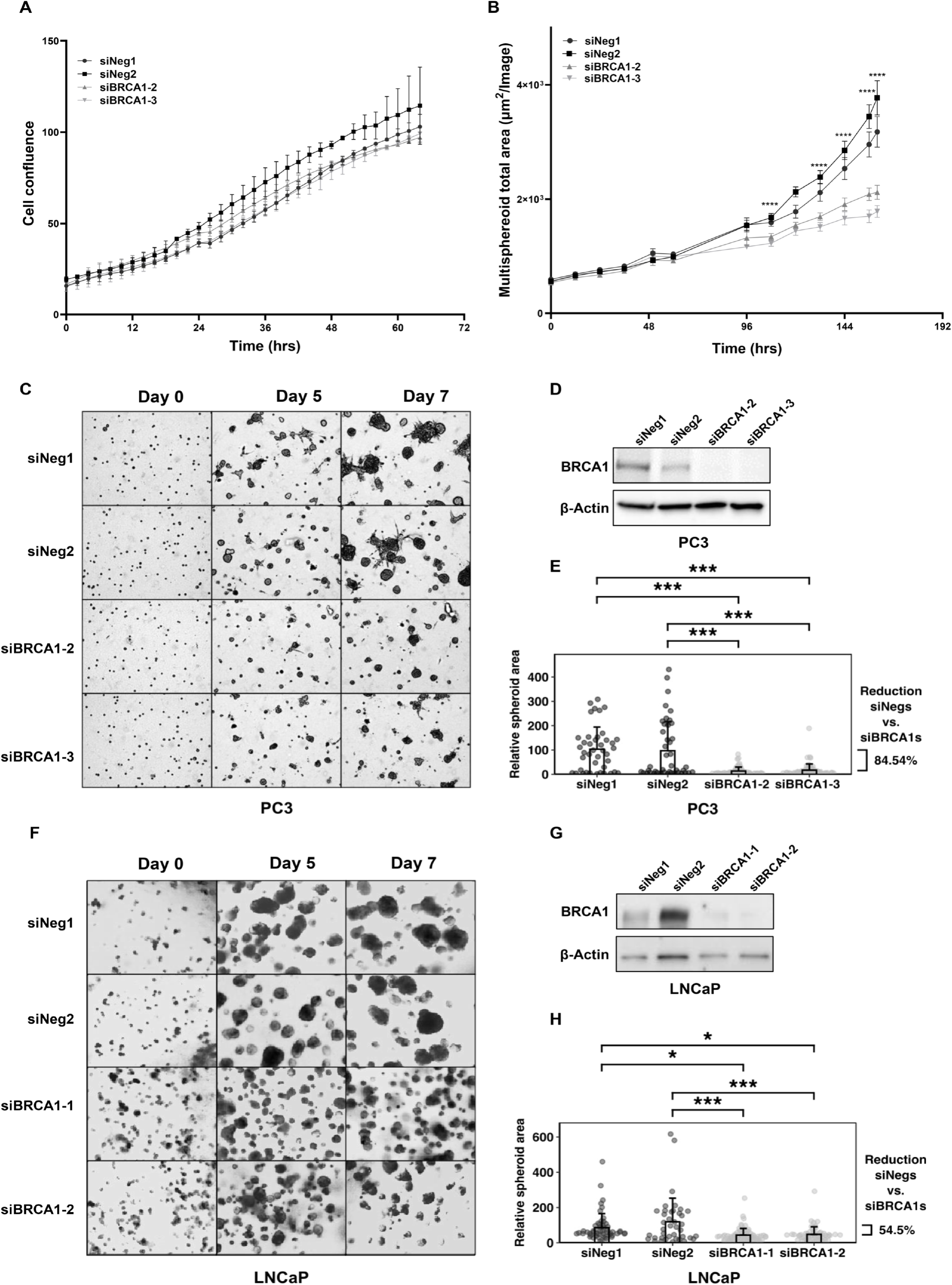
BRCA1 supports organotypic 3D spheroid growth of PCa cells. (A) Plot representing 2D proliferation with IncuCyte confluence analysis in PC3 cells 1–3 days after transfection with siNeg1, siNeg2, siBRCA1-2, or siBRCA1-3. (B) Plots representing 3D spheroid growth with IncuCyte analysis in PC3 cells 1–8 days after transfection with siNeg1, siNeg2, siBRCA1-2, or siBRCA1-3. (C) Representative brightfield images from days 0–7 in 3D culture of PC3 cells treated with siNeg1, siNeg2, siBRCA1-1, or siBRCA1-2. (D) Western blots depicting BRCA1 expression 48 h after transfection for 3D culture in PC3 cells. (E) Plots depicting relative 3D spheroid area in brightfield images for PC3 cells on day 7 of 3D culture. (F) Representative brightfield images from days 0–7 in a 3D culture of LNCaP cells. (G) Western blots depicting BRCA1 expression 48 h after transfection for 3D culture in LNCaP cells. (H) Plots depicting relative 3D spheroid area in brightfield images for LNCaP cells on day 7 of 3D culture.

### Disruption of mTOR complex suppresses HR repair while increasing sensitivity to AD in PCa

Given that BRCA1 interacts with and regulates the mTORC1 pathway, we evaluated the role of mTOR signaling in the regulation of DNA damage repair in PCa cells. VCaP cells were treated with the pan-mTOR inhibitor PP242 or vehicle in DNA damage –induced (cisplatin) and control cells. We analyzed the samples using Western blotting and observed that pan-mTOR inhibition modulated the response to cisplatin-induced DNA damage by inhibiting the induction of BRCA1 phosphorylation and Rad51 upregulation (Figure 6A). BRCA, Rad51, Cyclin A, and γH2AX were also visualized using immunofluorescence, where we observed upregulation, nuclear foci formation, and colocalization in response to cisplatin-induced DNA damage only when pan-mTOR inhibitor was not added, while γH2AX was still induced (Figure 6B, Supplementary Figure 5A and B). This suggested that the absence of functional mTOR signaling may prevent HR proteins from fully responding to cisplatin-induced DNA damage in PCa.

**Figure 6.**
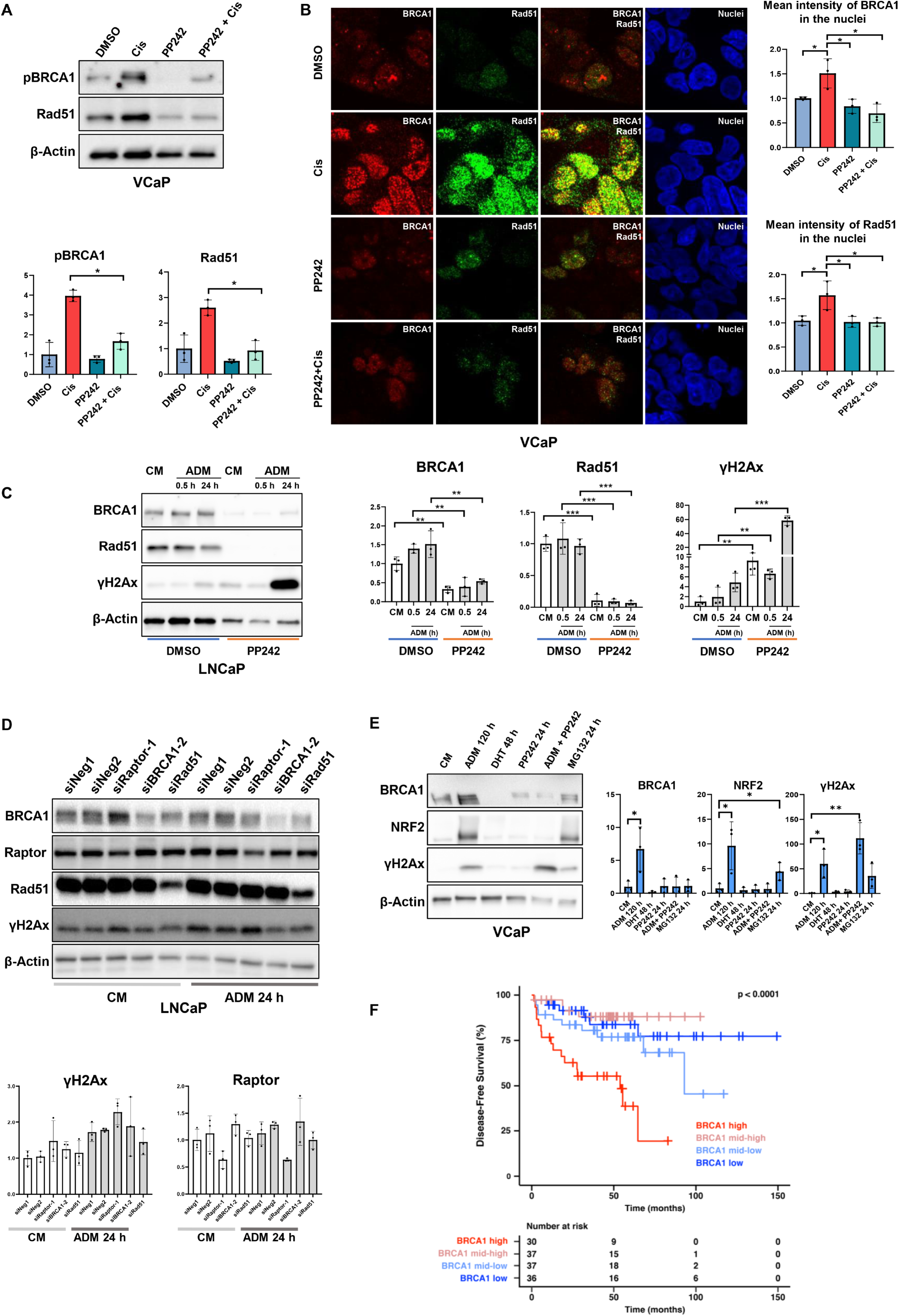
NRF2 regulation and DNA damage in response to AD are modulated by mTOR. (A) Western blots and bar graphs of pooled Western blot densitometry depicting pBRCA1, Rad51, and loading control expression after vehicle (DMSO) or cisplatin (Cis; 50 µM) in the presence or absence of pan-mTOR inhibitor PP242 (1 µM) in VCaP cells. (B) Representative immunofluorescence images of Rad51 (green) and BRCA1 (red) staining are represented along with merge images and bar graphs indicating mean intensities of Rad51 or BRCA1 in nuclei, after treatment with vehicle (DMSO) or cisplatin (50 µM) in the presence or absence of pan-mTOR inhibitor PP242 (1 µM) in VCaP cells. (C) Western blots and bar graphs of pooled Western blot densitometry depicting BRCA1, Rad51, γH2Ax, and loading control expression after vehicle (DMSO) or PP242 (1 µM) with or without ADM (0.5–24 h) in LNCaP cells. (D) Western blots and bar graphs of pooled Western blot densitometry depicting Raptor, γH2Ax, and loading control expression after transfection with siNeg1, siNeg2, siRaptor-1, siBRCA1-1 or siRad51 with or without ADM (24 h) in LNCaP cells. (E) Western blots and bar graphs of pooled Western blot densitometry depicting BRCA1, NRF2, γH2Ax, and loading control expression in VCaP cells subjected to vehicle (DMSO and/or methanol), 5 days of ADM, 48 h of DHT (10 nM) after 3 days of ADM, 24 h of PP242 (1 µM), 24 h of PP242 (1 µM) after 4 days of ADM, and 24 h of MG132 (10 µM) treatments. (F) Kaplan-Meier plot depicting disease-free survival of PCa patients by *BRCA1* mRNA quartiles in the MSK Cancer Cell 2010 cBioPortal dataset with significance determined using the log-rank test. The mean and standard deviation (SD) from three experiments are represented as bar graphs (*p < 0.05, **p < 0.01, and ***p < 0.001 as determined by *t-test*).

We tested whether inhibition of mTOR only inhibited HR function when DNA damage was induced using cisplatin. We induced DNA damage by depleting androgens in LNCaP cells, which we previously observed to induce γH2Ax and apoptosis (Figure 6A and B). In Western blot analysis, we observed that pan-mTOR inhibition enhanced the induction of DNA damage caused by AD in LNCaP cells (Figure 6C). Further, we tested whether disrupting the mTOR complex in PCa could enhance the induction of DNA damage. We transiently transfected LNCaP cells with Raptor siRNA to reduce the formation of the mTORC1 complex in the presence and absence of ADM. As expected, we observed a slight induction of γH2Ax upon Raptor knockdown (Figure 6D, Supplementary Figure S5C) suggesting that mTORC1 is required for the functioning of LNCaP and disrupting this complex would increase sensitivity towards ADM in PCa. This suggested that mTOR was also involved in regulating DNA repair in response to AD in PCa cells.

### AD upregulates antioxidant gene transcription factor NRF2

We previously observed that BRCA1 and γH2Ax levels are upregulated upon AD in PCa, based on our data from cell lines (Figure 2), xenografts, and PCa patients (Figure 3). This suggested that DDR might be active, even though Rad51 showed a trend of decrease, and the ATM response was inconsistent (Figure 2). Previous evidence had suggested that AD increases the expression of antioxidant markers and reactive oxygen species (ROS) in PCa cells, which is linked to castration resistance [32, 33]. Nuclear factor erythroid 2-related factor 2 (NRF2) is a well-attributed transcription factor that upregulates genes required for antioxidant synthesis [34]. BRCA1 is reported to regulate NRF2-dependent antioxidant signaling by physically interacting with NRF2 and promoting its stability and activation [35]. Here, we report that longer periods of AD (five days of ADM) upregulated NRF2 expression (Figure 6E, Supplementary Figure S5D). Furthermore, we also observe that ADM-induced NRF2 expression was blocked upon mTOR inhibition by PP242 along with BRCA1 (Supplementary figure S5D) suggesting that NRF2 expression is tightly controlled by BRCA1-mTOR signaling in PCa. Therefore, we suggest that upon AD, antioxidant synthesis may be activated by upregulated BRCA1 via NRF2 which could promote castration resistance in PCa.

Supporting the role of promoting aggressive features in PCa, high *BRCA1* expression was correlated with shorter disease-free and progression-free survival in patients (Figure 6F, Supplementary Figure S5E). While *BRCA1* is known to be mutated in PCa, amplified *BRCA1* is observed at similar rates in patients (Supplementary Figure S5F). Furthermore, neuroendocrine PCa (NEPC) features were more common in patients with high *BRCA1* expression levels (Supplementary Figure S4G). Together, these results suggest that BRCA1 may also be highly expressed in PCa, likely associated with cell survival thus promoting castration resistance.

## Discussion

Preclinical studies in PCa have shown that AR signaling regulates DDR and that, in turn, AR activation is enhanced upon DNA damage [9, 12]. Moreover, enzalutamide has been suggested to confer synthetic lethality in combination with PARPi by downregulating HR genes including BRCA1 [10, 11]. However, while improving overall survival of HRD patients in clinical trials, AR inhibition combined with PARPi has shown limited efficacy in HR-proficient patients [5, 7, 8, 36, 37]. This has prompted a re-examination of the current model of AR-DDR interplay, particularly in light of recent preclinical data suggesting that DDR regulation by AR is indeed not direct [13]. In addition, the low frequency of BRCA1 mutations in PCa compared to other HRD-prone cancer types remains unexplained [14]. Therefore, we sought to determine whether AR and BRCA1 have interplay in PCa and what the tissue-specific functions of BRCA1 in PCa are.

Our results demonstrate that ligand-activated AR strongly represses BRCA1 transcription, and thus, AD retains or even promotes BRCA1 expression. BRCA1 has been suggested to be repressed by Id4, the E2F family of transcription factors, Slug, and microRNAs [38–41]. This suggests that the AR regulates or interacts with these or other unknown repressors if it does not directly repress BRCA1. Moreover, we observed increased amount of activated BRCA1 in the post-ADT xenografts and patient samples. We further show that the decrease of BRCA1 expression by enzalutamide is utilizing AR-independent mechanisms. Therefore, it could be argued that the concept of enzalutamide-induced BRCAness/HRD suggested previously [10, 11] may not be restricted to AR-positive PCa and is mediated via unknown mechanisms. Enzalutamide has been previously shown also to downregulate RAD51C, RAD51AP1, RMI2, and RAD54L whose regulation by AR signaling was not examined in our study [10]. Nevertheless, events activating BRCA1 expression after AD may counteract the previously suggested enzalutamide-induced BRCAness/HRD and its synergy with PARPi.

Although the key HR components, BRCA1 and Rad51, were strongly downregulated by AR activation, no DNA damage was observed. Interestingly, conversely, ADM induced DNA damage in all tested cell lines while simultaneously increasing the expression or phosphorylation of BRCA1. We further examined the response to AD by studying the acute effects of ADT on castrated VCaP xenografts. While ADM is devoid of steroids, castration/ADT specifically reduces androgens; therefore, observing similar results in ADM experiments and the murine model suggested that results obtained from cell lines were indeed related to AD. Moreover, by studying patient samples before and after ADT we observed an increase in both the expression and phosphorylation of BRCA1 and sustained DNA damage. The 22Rv1 cell line highly expresses AR-V7, but also expresses full-length AR. Thus, DNA damage in ADM is not a consequence of the absence of AR signaling but, more specifically, the absence of full-length AR signaling. Full-length AR contains the ligand-binding domain which also interacts with coregulators and is used in forming a complex with nuclear mTOR (nmTOR) [42]. Active AR may bind DNA together with nmTOR [43]. Thus, ADT facilitates the release of AR-bound nmTOR which can then be observed to transiently increase pS6 in PTEN intact cell lines. Once the AR-nmTOR complex is disrupted it no longer sustains the transcription and expression of mTOR, contributing to the delayed loss of mTOR signaling in response to ADM. Our results demonstrate that the loss of HR-related gene expression in response to AR activation is not sufficient to induce DNA damage by itself. Moreover, BRCA1 activation during acute AD cannot prevent DNA damage. Instead, mTOR signaling appears to play a more relevant role in AD-induced DNA damage, and thus, BRCA1 may contribute to AD predominantly via its interaction with Raptor.

Our results demonstrate a novel interplay between BRCA1 and mTORC1 in PCa via Raptor based on an unbiased mass spectrometry pulldown assay and verification by IP Western blot and immunofluorescence using multiple antibodies. We also demonstrate a functional connection between BRCA1 and mTOR by showing that during BRCA1 knockdown, S6 phosphorylation is reduced. The acute response to stress induced by AD includes a surge in the phosphorylation of PI3K-Akt/mTOR pathway components. Our results show that BRCA1 downregulation reduces S6 phosphorylation, suggesting that high BRCA1 protein levels could influence cancer cell survival during AD by potentiating the initial surge of survival signaling mediated by S6. Furthermore, our data demonstrate that the mTOR inhibitor PP242 downregulates the expression of the HR genes BRCA1 and Rad51. PP242 disrupts the partaking of BRCA1 and Rad51 in DNA repair complexes after cisplatin treatment and severely enhances DNA damage in response to AD. However, transient knockdown reveals that the mTOR complex itself has a greater protective effect on DNA during AD than BRCA1 or Rad51 alone. This suggests that the role of HR is lesser than that of mTOR in AD-induced DNA damage and the role of HR may contribute via the interaction between BRCA1 and Raptor/mTORC1. It might also be that some of the previously AR-contributed regulation of DDR is in fact mTOR-mediated.

We demonstrated that AD induces DNA damage, followed by upregulation of BRCA1 and NRF2. AD is known to induce ROS-mediated oxidative stress and BRCA1 regulates NRF2 [32–35]. NRF2 was strongly upregulated after five days of AD, suggesting that NRF2 expression might have been too low to be detected as a significant BRCA1 pull-down hit in mass spectrometry under normal culture conditions. Low NRF2 levels have been associated with promoting PCa initiation by increasing ROS, but NRF2 expression is higher in CRPC than in primary tumors [44, 45]. Therefore, the cessation of BRCA1 suppression by AR during AD may work as a protective mechanism against oxidative stress-induced DNA damage by initiating NRF2-mediated antioxidative processes via BRCA1 upregulation.

We observed that 3D growth was reduced upon BRCA1 knockdown in AR-positive and AR-negative cell lines. Patient data also suggested that high *BRCA1* expression was linked to shorter disease-free survival and that *BRCA1* was amplified in PCa. This provides evidence that BRCA1 functions beyond its the role as a tumor suppressor in PCa. BRCA1 may promote organotypic growth by interacting with mTOR signaling or by other functions suggested by the novel interactors. High BRCA1 levels are also associated with neuroendocrine PCa features. Interestingly, a recent patient-derived organoid study showed increased DDR gene expression in neuroendocrine samples, which might suggest that AR independence is linked to the increased function of some DDR genes [46].

In conclusion, our study sheds light on the multifaceted role of BRCA1 in PCa, particularly its novel interaction with the mTORC1 pathway and its role in cell growth. We demonstrate that ADT activates rather than suppresses BRCA1 expression in PCa. Thus, we conclude that BRCA1 plays a dual role in PCa, acting as a tumor suppressor when mutated but also paradoxically promoting cancer in PCa driven by different biological developmental paths.

## Acknowledgements

Mass spectrometry analysis was performed with the help of Mirva Pääkkönen at the Turku Proteomics Facility, University of Turku and Åbo Akademi University. The facility is supported by Biocenter Finland. We thank Sinikka Collanus and the Histology core facility of the Institute of Biomedicine, University of Turku, for their assistance with IHC. We thank Minna Santanen, Jukka Karhu, and Barbara Ramos Artigot for their excellent technical assistance. We also thank Turku Center for Disease Modeling (TCDM), Guillermo Martinez Nieto, and Petra Sipilä for providing their expertise in generating mouse xenograft models. We also express our gratitude to Dr. Jouko Sandholm and Cell Imaging and Cytometry Core for providing consultation regarding microscopy and image analysis.

## Author Contributions statement

SS, VV, and MS designed the study, conceived the experiments, and analyzed the data. SS and VV prepared figures. SS, VV, AK, AL, and MT carried out experiments. SS, VV, AK, and MS designed in silico analyses. JA and AR performed the mass spectrometry pulldown experiments. MP designed the xenograft experiment. PT designed the human immunohistochemistry analyses and provided the samples. GW analyzed the human immunohistochemistry samples. SS, VV, and MS wrote the manuscript and all authors commented on and approved the final version. MS acquired the funding and supervised the study.

