## Supplementary_Figures for "Functionality of BRCA1 supports the survival of prostate cancer cells during the development of castration resistance"

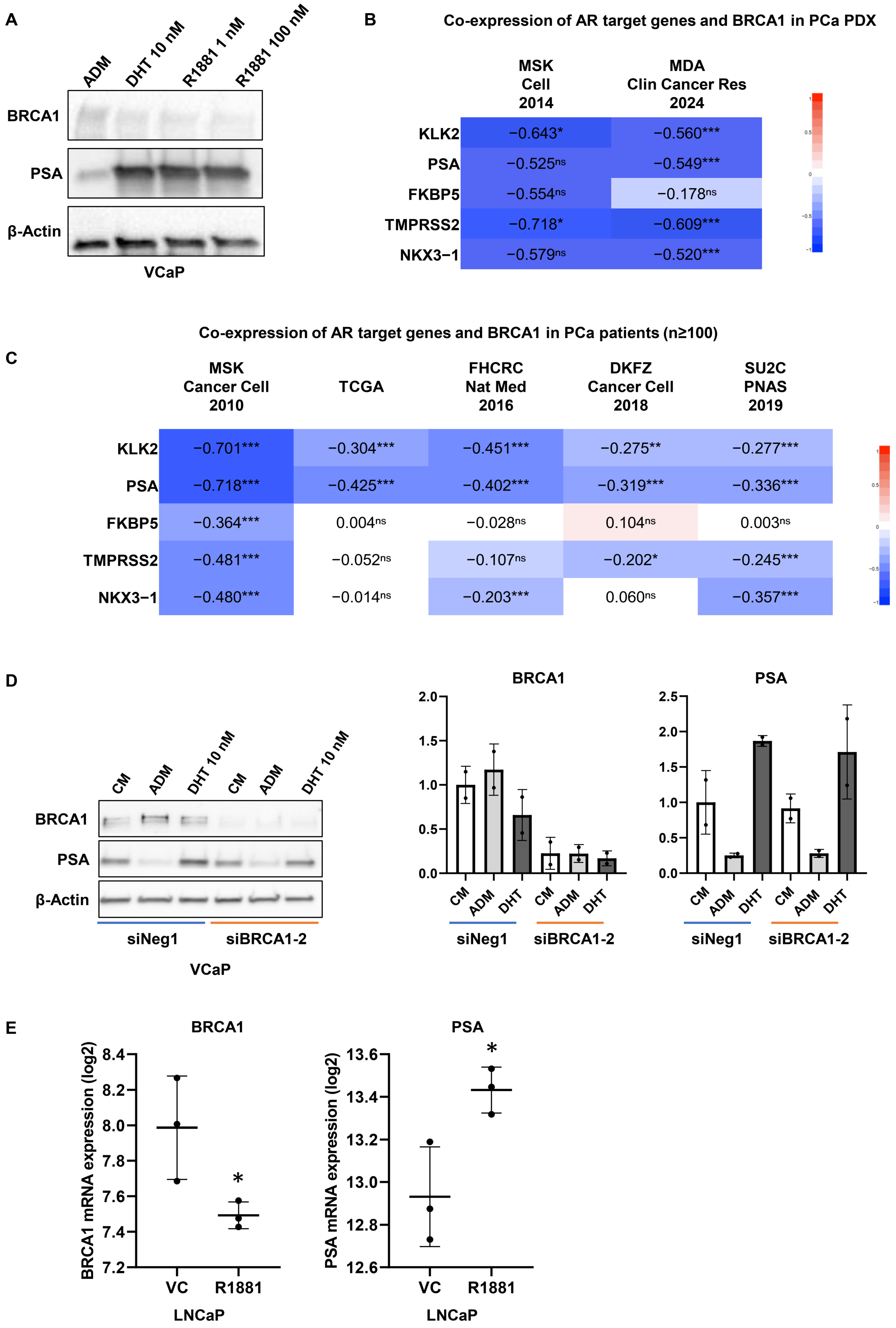
A
ADM
DHT 10 nM
R1881 1 nM
R1881 100 nM
BRCA1
PSA
β-Actin
VCaP
B
Co-expression of AR target genes and BRCA1 in PCa PDX
MSK Cell 2014
MDA Clin Cancer Res 2024
KLK2 −0.643* −0.560***
PSA −0.525ns −0.549***
FKBP5 −0.554ns −0.178ns
TMPRSS2 −0.718* −0.609***
NKX3−1 −0.579ns −0.520***
Co-expression of AR target genes and BRCA1 in PCa patients (n≥100)
C
MSK Cancer Cell 2010
TCGA
FHCRC Nat Med 2016
DKFZ Cancer Cell 2018
SU2C PNAS 2019
KLK2 −0.701*** −0.304*** −0.451*** −0.275** −0.277***
PSA −0.718*** −0.425*** −0.402*** −0.319*** −0.336***
FKBP5 −0.364*** 0.004ns −0.028ns 0.104ns 0.003ns
TMPRSS2 −0.481*** −0.052ns −0.107ns −0.202* −0.245***
NKX3−1 −0.480*** −0.014ns −0.203*** 0.060ns −0.357***
D
CM
ADM
DHT 10 nM
BRCA1
PSA
β-Actin
siNeg1
siBRCA1-2
VCaP
BRCA1
PSA
CM
ADM
DHT
E
BRCA1
PSA
BRCA1 mRNA expression (log2)
PSA mRNA expression (log2)
VC
R1881
LNCaP

**Supplementary figure S1. Association between AR and BRCA1 in androgen-dependent PCa.**

(A) Representative Western blots of BRCA1, PSA, and loading control expression in VCaP cells subjected to 72 h of ADM, 24 h of DHT (10 nM), and 24 h R1881 (1 nM and 100 nM) treatments following 48 h of ADM. Vehicle (methanol and/or ethanol) were added to wells not containing DHT and/or R1881 respectively. (B) Heat maps representing co-expression between selected AR target genes and *BRCA1* in PCa patient datasets available at cBioPortal with $n \geq 100$ (ns = not significant, $^{*}p < 0.05$, and $^{***}p < 0.001$ as determined for the significance level of Pearson coefficient). (C) Heat maps representing co-expression of AR target genes and *BRCA1* in PCa PDX datasets available at cBioPortal (ns = not significant, $^{*}p < 0.05$, $^{**}p < 0.01$ and $^{***}p < 0.001$ as determined for the significance level of Pearson coefficient). (D) Western blots and bar graphs of pooled Western blot densitometry depicting BRCA1, PSA, and loading control expression after 72 h of ADM and 24 h of DHT following 48 h of ADM treatments with or without silencing BRCA1 in VCaP cells. (E) Graphs representing *BRCA1* and *PSA* mRNA expression levels in LNCaP cells subjected to vehicle (DMSO) or R1881 treatment, analyzed from the GSE50936 dataset ($^{*}p < 0.05$ as determined by *t-test*).

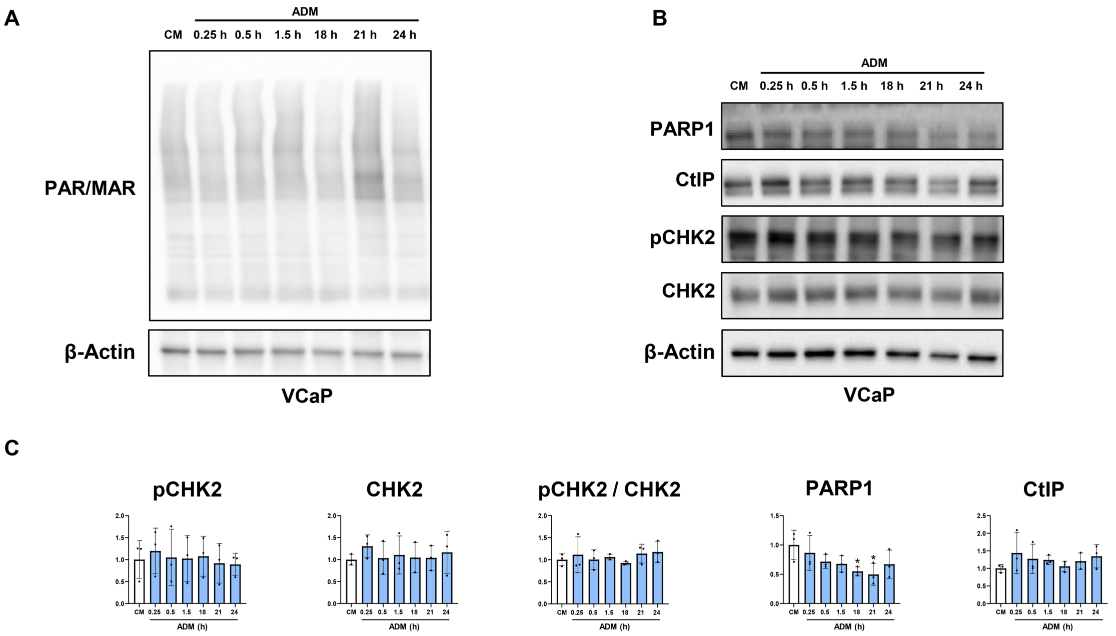


**Supplementary figure S2. Analysis of additional DDR markers in AD conditions and influence of mTOR signaling pathway in HR repair pathway in PCa.**

(A-C) Representative Western blots and bar graphs of pooled Western blot densitometry depicting additional markers of DSB repair (CtIP, pCHK, and CHK2), SSB repair (PARP1 and PAR/MAR), and loading control in VCaP cells subjected to ADM (0.25–24 h) or control media. The mean and standard deviation (SD) from three experiments are represented as bar graphs.

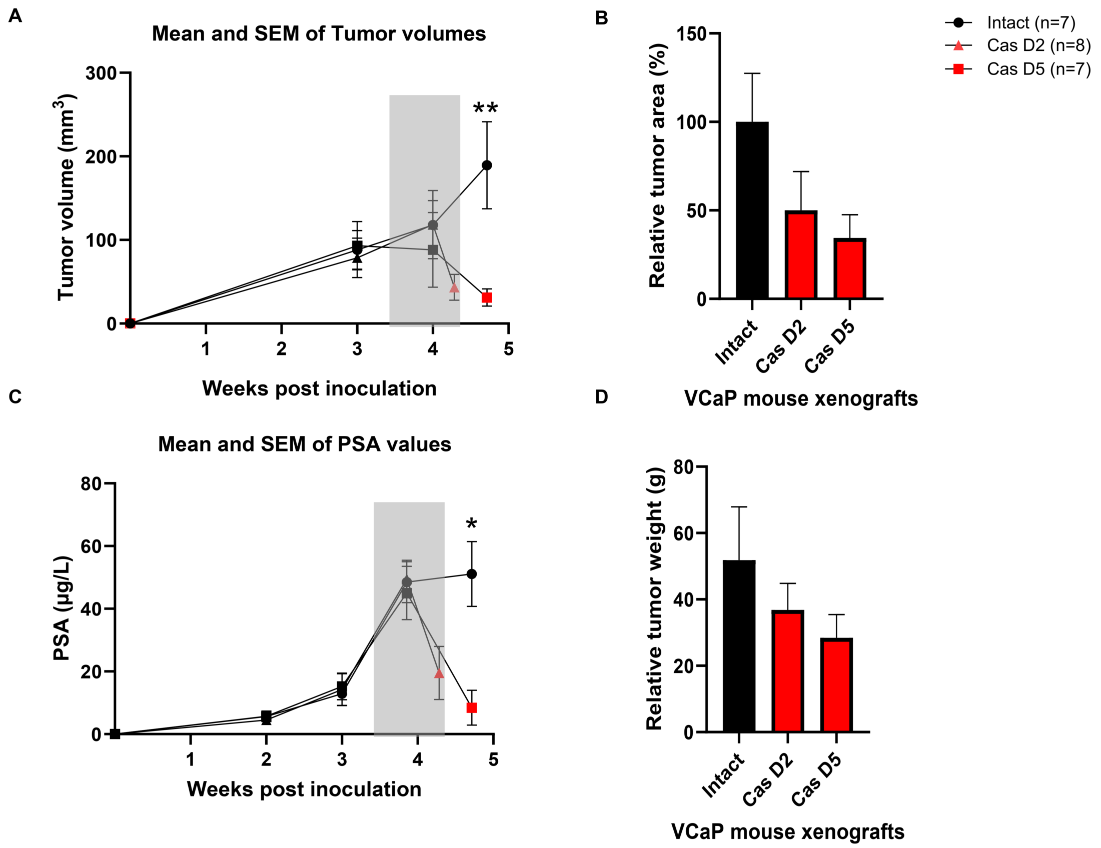


**Supplementary figure S3. Representation of key attributes of Intact, Cas D2, and Cas D5 tumors obtained from VCaP mice xenografts.**

(A) Graphs representing mean and SEM of tumor volumes ($mm^3$) of Intact, Cas D2, and Cas D5 xenografts (**$p < 0.01$ as determined by the *t-test*). (B) Bar graphs representing relative tumor area measured from Intact, Cas D2, and Cas D5 xenograft tumor sections. (C) Graphs representing mean and SEM of PSA (µg/l) values for Intact, Cas D2, and Cas D5 mice (*$p < 0.01$ as determined by the *t-test*). (D) Bar graphs representing relative tumor weight (g) of Intact, Cas D2 and Cas D5 tumors obtained from VCaP mice xenografts.

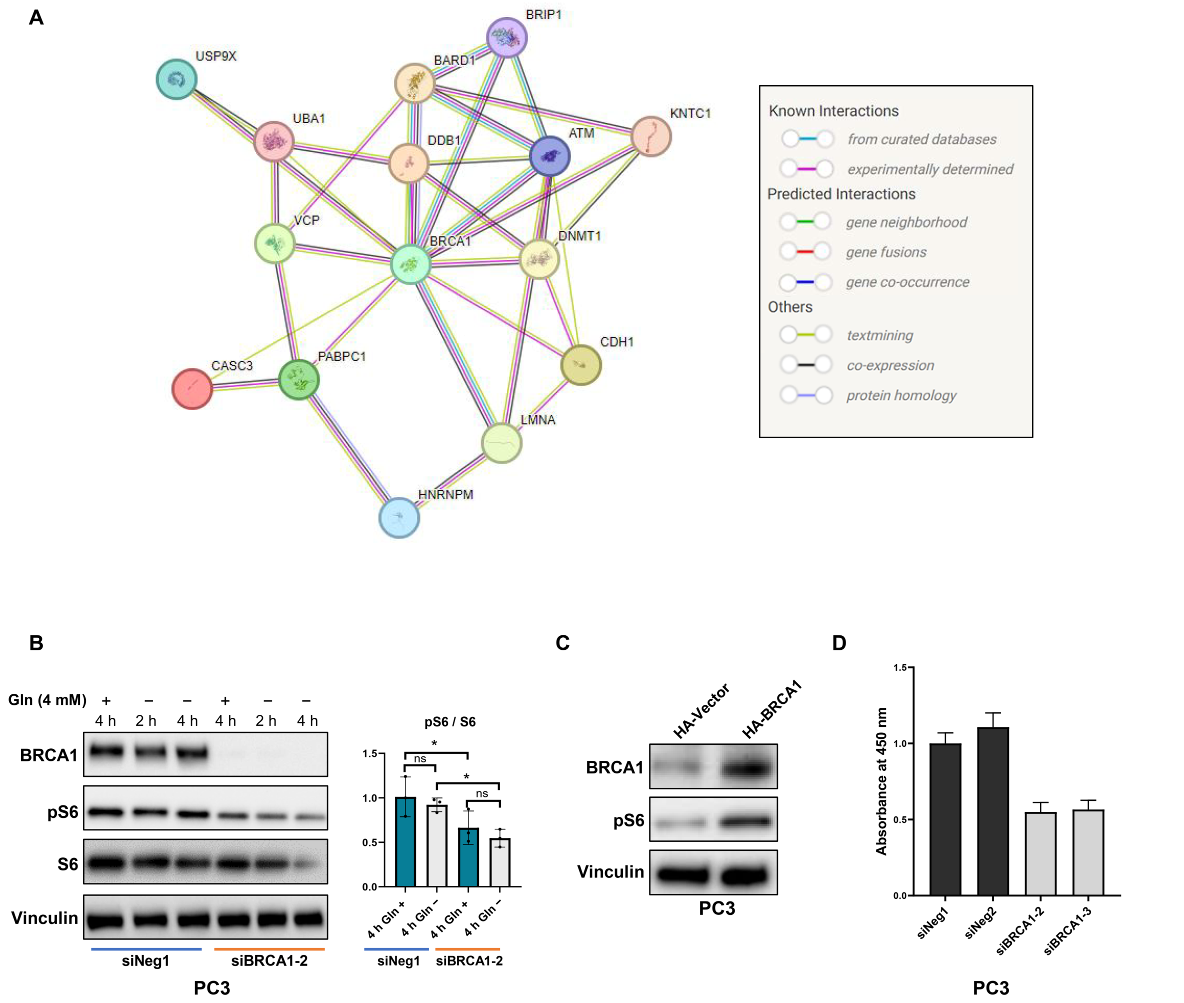


**Supplementary figure S4. Association of BRCA1 and mTOR signaling pathway in PCa.** (A) StringDB analysis depicting existing evidence for interactions between BRCA1 and identified mass spectrometry hits. (B) Representative Western blot and bar graphs of pooled Western blot densitometry depicting BRCA1, pS6, S6, and loading control expression after transfection with siNeg1 or siBRCA1-2 and glutamine deprivation for 2–4 h preceding lysis in PC3 cells (ns = not significant and *$p < 0.05$ as determined by the *t-test*). (C) Western blot images of pS6, BRCA1, and loading control expression in PC3 cells overexpressing HA-empty vector or HA-tagged BRCA1 proteins achieved by generating stable cell line. (D) Bar graphs representing viability of PC3 spheroids after silencing BRCA1 as determined by WST-8 assay at day 7 of growth.

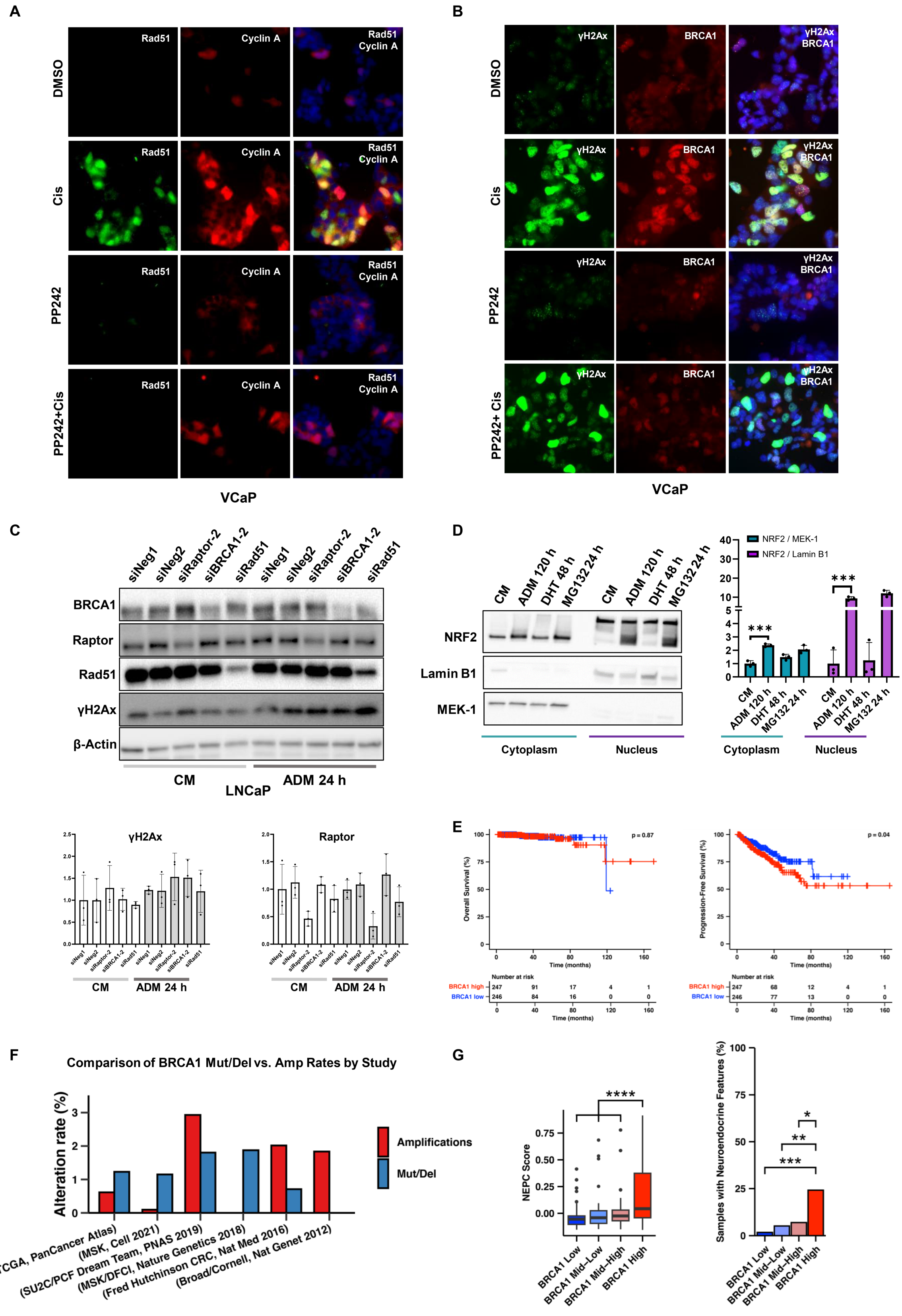
A
Rad51
Cyclin A
Rad51 Cyclin A
DMSO
Cis
PP242
PP242+Cis
VCaP
B
γH2Ax
BRCA1
γH2Ax BRCA1
DMSO
Cis
PP242
PP242+ Cis
VCaP
C
siNeg1 siNeg2 siRaptor-2 siBRCA1-2 siRad51 siNeg1 siNeg2 siRaptor-2 siBRCA1-2 siRad51
BRCA1
Raptor
Rad51
γH2Ax
β-Actin
CM
ADM 24 h
LNCaP
γH2Ax
Raptor
CM
ADM 24 h
D
CM ADM 120 h DHT 48 h MG132 24 h CM ADM 120 h DHT 48 h MG132 24 h
NRF2
Lamin B1
MEK-1
Cytoplasm
Nucleus
NRF2 / MEK-1
NRF2 / Lamin B1
***
***
Cytoplasm
Nucleus
E
p = 0.87
Overall Survival (%)
Time (months)
Number at risk
BRCA1 high 247 91 17 4 1
BRCA1 low 246 84 16 0 0
p = 0.04
Progression-Free Survival (%)
Time (months)
Number at risk
BRCA1 high 247 68 12 4 1
BRCA1 low 246 77 13 0 0
F
Comparison of BRCA1 Mut/Del vs. Amp Rates by Study
Alteration rate (%)
Amplifications
Mut/Del
(TCGA, PanCancer Atlas)
(MSK, Cell 2021)
(SU2C/PCF Dream Team, PNAS 2019)
(MSK/DFCI, Nature Genetics 2018)
(Fred Hutchinson CRC, Nat Med 2016)
(Broad/Cornell, Nat Genet 2012)
G
NEPC Score
****
BRCA1 Low
BRCA1 Mid-Low
BRCA1 Mid-High
BRCA1 High
Samples with Neuroendocrine Features (%)
***
**
*

**Supplementary figure S5. Association between DNA damage, AD, and mTOR signaling pathway in PCa.**

(A) Representative immunofluorescence images of Rad51 (green), Cyclin A (red), and merge after treatment with vehicle (DMSO) or cisplatin (50 µM) in the presence or absence of pan-mTOR inhibitor PP242 (1 µM) in VCaP cells. (B) Representative immunofluorescence images of γH2Ax (green), BRCA1 (red), and merge after treatment with vehicle (DMSO) or cisplatin (50 µM) in the presence or absence of pan-mTOR inhibitor PP242 (1 µM) in VCaP cells. (C) Representative Western blots and bar graphs of pooled Western blot densitometry representing BRCA1, Raptor, Rad51, γH2Ax, and loading control expression in cells subjected to silencing of Raptor, Rad51 or BRCA1 and control media or 24 h of ADM in LNCaP cells. (D) Western blots and bar graphs of pooled Western blot densitometry depicting NRF2, Lamin B1 (Nuclear), and MEK-1 (Cytoplasm) expression in cytoplasmic and nuclear fractions of VCaP cells subjected to vehicle (DMSO and/or methanol), 5 days of ADM, 48 h of DHT (10 nM) after 3 days of ADM, and 24 h of MG132 (10 µM) treatments preceding lysis. (E) Kaplan-Meier plot depicting overall survival and progression-free survival of PCa patients by *BRCA1* mRNA medians in the TCGA PanCancer Atlas cBioPortal dataset with significance determined using the log-rank test. (F) Plots representing BRCA1 alterations in PCa Patient datasets obtained from cBioPortal. (G) Plots representing the presence of histological NEPC features and transcript-based NEPC score in PCa patients by *BRCA1* mRNA expression quartiles in the SU2C PNAS 2019 cBioPortal dataset.
