## Supplementary_Tables for "Functionality of BRCA1 supports the survival of prostate cancer cells during the development of castration resistance"

**Supplementary Table 1. Antibodies.** pAb, polyclonal antibody; mAb, monoclonal antibody; Xe, xenografts; Pa, patient samples.

| Antigen | Cat.no | Type | Dilution | | | | Source |
|---|---|---|---|---|---|---|---|
| | | | WB | IHC | IP/ MS | IF | |
| β-Actin | sc-47778 | Mouse mAb | 1:1000 | - | - | - | Santa Cruz Biotechnology |
| AKT (Pan) | CST#4691 | Rabbit mAb | 1:1000 | - | - | - | Cell Signaling Technology |
| pAKT (s473) | CST#4060 | Rabbit mAb | 1:1000 | - | - | - | Cell Signaling Technology |
| AR | sc-816 | Rabbit mAb | 1:1000 | - | - | - | Santa Cruz Biotechnology |
| ASH2L | CST#5019 | Rabbit mAb | 1:1000 | - | - | - | Cell Signaling Technology |
| ATM | sc-377293 | Mouse mAb | 1:1000 | - | - | - | Santa Cruz Biotechnology |
| pATM (s1981) | CST#5883 | Rabbit mAb | 1:1000 | - | - | - | Cell Signaling Technology |
| BRCA1 | sc-6954 | Mouse mAb | - | - | ✓ | 1:50 | Santa Cruz Biotechnology |
| BRCA1 | CST#9010 | Rabbit mAb | 1:1000 | - | - | 1:200 | Cell Signaling Technology |
| BRCA1 | MS110 | Mouse mAb | - | 1:50 | - | - | Millipore Corporation |
| pBRCA1 (Ser1524) | CST#9009 | Rabbit mAb | 1:1000 | - | - | - | Cell Signaling Technology |
| pBRCA1 (Ser1423) | ab47325 | Rabbit pAb | - | 1:500 (Xe)<br>1:150 (Pa) | - | - | Abcam |
| CHK2 | CST#2662 | Rabbit mAb | 1:1000 | - | - | - | Cell Signaling Technology |
| pCHK2 (t68) | CST#2197 | Rabbit mAb | 1:1000 | - | - | - | Cell Signaling Technology |
| Cleaved-Caspase-3 | CST#9664 | Rabbit mAb | 1:1000 | - | - | - | Cell Signaling Technology |
| CtIP | sc-271339 | Mouse mAb | 1:1000 | - | - | - | Santa Cruz Biotechnology |
| Cyclin-A | sc-271682 | Mouse mAb | 1:1000 | - | - | 1:200 | Santa Cruz Biotechnology |
| FKBP5 | CST#12210 | Rabbit mAb | 1:1000 | - | - | - | Cell Signaling Technology |
| Phospho-Histone H2A.X (Ser139) | CST#9718 | Rabbit mAb | 1:1000 | 1:200 | - | 1:200 | Cell Signaling Technology |
| Lamin-B1 | sc-377000 | Mouse mAb | 1:1000 | - | - | - | Santa Cruz Biotechnology |
| MEK-1 | sc-6250 | Mouse mAb | 1:1000 | - | - | - | Santa Cruz Biotechnology |
| NRF2 | ab62352 | Rabbit mAb | 1:1000 | - | - | - | Abcam |
| PAR/MAR | CST#83732 | Rabbit mAb | 1:1000 | - | - | - | Cell Signaling Technology |

| | | | | | | | |
|---|---|---|---|---|---|---|---|
| PARP1 | sc-8007 | Mouse mAb | 1:1000 | - | - | - | Santa Cruz Biotechnology |
| PSA/KLK3 | CST#5365 | Rabbit mAb | 1:1000 | - | - | - | Cell Signaling Technology |
| Rad51 | ab133534 | Rabbit mAb | - | 1:500 | - | - | Abcam |
| Rad51 | CST#8875 | Rabbit mAb | 1:1000 | - | - | 1:200 | Cell Signaling Technology |
| Raptor | sc-518004 | Mouse mAb | 1:1000 | - | - | 1:100 | Santa Cruz Biotechnology |
| Raptor | CST#2280 | Rabbit mAb | 1:1000 | - | - | 1:100 | Cell Signaling Technology |
| Raptor | MP-05-1470 | Mouse mAb | 1:1000 | - | - | 1:100 | Millipore Corporation |
| RBBP5 | CST#13171 | Rabbit mAb | 1:1000 | - | - | - | Cell Signaling Technology |
| S6 | CST#2317 | Mouse mAb | 1:500 | - | - | - | Cell Signaling Technology |
| pS6 (Ser240/244) | CST#5364 | Rabbit mAb | 1:1000 | 1:1000 | - | - | Cell Signaling Technology |
| Vinculin | sc-73614 | Mouse mAb | 1:1000 | - | - | - | Santa Cruz Biotechnology |
| Peroxidase-conjugated goat anti-rabbit IgG | A16104 | Rabbit pAb | 1:10000 | - | - | - | Invitrogen |
| Peroxidase-conjugated goat anti- mouse IgG | A16072 | Mouse pAb | 1:10000 | - | - | - | Invitrogen |
| Alexa Fluor Anti-rabbit 488 nm | A-11034 | Rabbit pAb | - | - | - | 1:1000 | Invitrogen |
| Alexa Fluor Anti-mouse 555 | A-28180 | Mouse pAb | - | - | - | 1:1000 | Invitrogen |
| IgG mouse | sc-2025 | normal mouse IgG | - | - | ✓ | - | Santa Cruz Biotechnology |

**Supplementary Table 2. siRNAs.**

| siRNA | Cat no | Target gene | Concentration (nM) | | |
|---|---|---|---|---|---|
| | | | PC3 | VCaP | LNCaP |
| siBRCA1-1 | siRNA#457 | *BRCA1* | 2.5 | 10 | 5 |
| siBRCA1-2 | siRNA#458 | *BRCA1* | 2.5 | 10 | 5 |
| siBRCA1-3 | siRNA#459 | *BRCA1* | 2.5 | 10 | 5 |
| siRaptor-1 | siRNA#33214 | *Raptor* | 2.5 | 10 | 5 |
| siRaptor-2 | siRNA#33215 | *Raptor* | 2.5 | 10 | 5 |
| siRad51 | siRNA#11735 | *Rad51* | 2.5 | 10 | 5 |
| siNeg1 | Neg CTRL#1 | - | 2.5 | 10 | 5 |
| siNeg2 | Neg CTRL#2 | - | 2.5 | 10 | 5 |
